# *CREBBP* and *STAT6* co-mutation and 16p13 and 1p36 loss define the t(14;18)-negative diffuse variant of follicular lymphoma

**DOI:** 10.1101/2020.05.28.120212

**Authors:** Rena R. Xian, Yi Xie, Lisa M. Haley, Raluca Yonescu, Aparna Pallavajjala, Stefania Pittaluga, Elaine S. Jaffe, Amy S. Duffield, Chad M. McCall, Shereen M. F. Gheith, Christopher D. Gocke

## Abstract

The diffuse variant of follicular lymphoma (dFL) is a rare variant of FL lacking t(14;18) that was first described in 2009. In this study, we use a comprehensive approach to define unifying pathologic and genetic features through gold-standard pathologic review, FISH, SNP-microarray and next-generation sequencing of 16 cases of dFL. We found unique morphologic features, including interstitial sclerosis, microfollicle formation, and rounded nuclear cytology, confirmed absence of t(14;18) and recurrent deletion of 1p36, and showed a novel association with deletion/CN-LOH of 16p13 (inclusive of *CREBBP, CIITA* and *SOCS1*). Mutational profiling demonstrated near-uniform mutations in *CREBBP* and *STAT6*, with clonal dominance of *CREBBP*, among other mutations typical of germinal-center B-cell lymphomas. Frequent *CREBBP* and *CIITA* co-deletion/mutation suggested a mechanism for immune evasion, while subclonal *STAT6* activating mutations with concurrent *SOCS1* loss suggested a mechanism of BCL-xL/BCL2L1 upregulation in the absence of *BCL2* rearrangements. A review of the literature showed significant enrichment for 16p13 and 1p36 loss/CN-LOH, *STAT6* mutation, and *CREBBP* and *STAT6* co-mutation in dFL, as compared to conventional FL. With this comprehensive approach, our study demonstrates confirmatory and novel genetic associations that can aid in the diagnosis and subclassification of this rare type of lymphoma.

## Introduction

Follicular lymphoma (FL) is the second most common nodal non-Hodgkin lymphoma accounting for approximately 20% of all lymphomas (1). The proliferation of germinal center B-cells (GCB) forming abnormal follicles coupled with translocation of the anti-apoptotic gene *BCL2* with *IGH* resulting in t(14;18)(q32;q21) are diagnostic hallmarks of FL (1). However, there are exceptions, as approximately 5% of low-grade follicular lymphoma (LGFL) show a predominantly diffuse growth pattern (2, 3), and approximately 10% of FL lack t(14;18) (1), most of which represent high-grade disease.

The 2016 WHO classification recognizes several variants and related entities of FL, the latter of which is designated as conventional follicular lymphoma (cFL). The morphologically low-grade spectrum includes in-situ follicular neoplasia, duodenal-type FL, and the diffuse FL variant (dFL) with the former two entities consistently demonstrating t(14;18) *BCL2/IGH* rearrangements. The morphologically high-grade spectrum includes testicular FL and pediatric-type FL (pFL), neither of which carry *BCL2/IGH* rearrangements. Genomic analysis of cFL has shown that in addition to t(14;18), a number of recurrent copy number variants (CNVs) (4-10) and somatic mutations can be found (11-19), such as CNVs of 1p36, mutations of epigenetic regulators *KMT2D, CREBBP* and *EZH2*, and mutations of *TNFRSF14*. The genetic abnormalities found in cFL serve as the basis against which variant subtypes can be compared.

dFL is the only LGFL variant lacking t(14;18). This entity was first described in 2009 in 35 cases as an unusual type of LGFL with a predominantly diffuse growth pattern, characteristic immunophenotype, and near-uniform deletion of chromosome 1p36 (3). This variant of FL was distinguished from LGFL with a predominantly diffuse growth pattern, as the former consistently lack the characteristic *BCL2* rearrangement, whereas the latter consistently demonstrate t(14;18). Besides the genetic difference, the 2009 description of dFL also found characteristic clinical features, such as frequent groin/inguinal site of presentation, bulky low clinical stage disease, and good prognosis. Subsequent to this description, two other series evaluating 11 cases (20) and 6 cases (21) of dFL confirmed recurrent 1p36 abnormalities and/or *TNFRSF14* mutations (20), as well as mutations of *CREBBP* and *STAT6* (20, 21). The aims of the present study was to use a comprehensive approach to build upon existing, yet incomplete, literature, and determine unifying pathologic and genetic abnormalities. Findings from this study improve our understanding of the relationship between dFL and cFL, the molecular pathogenesis of dFL, and identifies potential molecular markers that may aid in the diagnosis and accurate subclassification of this rare variant of FL.

## Methods

### Pathologic Case Selection

Excisional biopsies were selected from the pathology archives of the Johns Hopkins Hospital (JHH) and the National Cancer Institute (NCI) after appropriate institutional review board approval. The JHH archives were searched from 1984-2013 for all cases of LGFL with a predominantly diffuse growth pattern (≥75%), occurring in an inguinal/groin site, and showing co-expression of CD23. The NCI archives were searched from 2000-2014 for cases citing the original Katzenberger *et al* description (3). BCL2 protein expression by immunohistochemistry and BCL2 rearrangement status for FISH were not used as selection criteria. Cases with components of histologic grade 3 or diffuse large B-cell lymphoma (DLBCL), or with high proliferation indices (>30%) were excluded. The histologic and immunohistochemical stains, and available clinical and ancillary data, were reviewed in concert by the study authors with consensus agreement of the final diagnoses.

### Fluorescence In-Situ Hybridization

FISH using the Vysis LSI *IGH/BCL2* dual color dual fusion probe (Abbott Molecular, Des Plaines, IL) was performed on all cases (per manufacturer’s protocol) without clinically available FISH analysis using formalin-fixed paraffin embedded (FFPE) tissue. Since histologic sectioning results in overlapping cells, only dual fusion signals identified in the same plane were considered as true fusions. Detailed methods can be found in *Supplementary Information*.

### SNP-Microarray Analysis

DNA was extracted using 4-10 unstained slides of FFPE tissue using the Pinpoint Slide DNA Isolation system (Zymo Research, Orange, CA). In brief, unstained slides were deparaffinized with xylene followed by tissue dissection and Proteinase K digestion. DNA cleanup was performed using the QIAamp DNA mini kit with the QIAcube instrument (QIAGEN, Valencia, CA). SNP-microarray using the HumanCytoSNP12 BeadChip platform (Illumina, San Diego, CA), which assesses approximately 300,000 polymorphic loci, was performed per manufacturer’s protocol. Analysis was completed using KaryoStudio (Illumina, San Diego, CA) and Nexus Copy Number (BioDiscovery, Hawthorne, CA). SNP and gene annotations were compared against the National Center for Biotechnology Information (NCBI) genome build 37 (GRCh37/hg19). CNVs were determined by independent and consensus review by R.R.X and C.D.G. Detailed interpretive criteria can be found in *Supplementary Information.*

### Targeted Next-Generation Sequencing Analysis

Using 200 ng of extracted DNA from the above analysis, DNA hybrid capture libraries were prepared using an Agilent SureSelect-XT (Agilent Technologies, Santa Clara, CA) custom-designed target enrichment kit evaluating full-gene sequences of 641 cancer-related genes (see *Supplementary Information*), as previously described (22). Following DNA quality control, shearing and library preparation, next generation sequencing was performed on an Illumina HiSeq 2500 (Illumina Biotechnology, San Diego, CA) using 2×100bp Rapid Run v2 paired end chemistry. Using vendor supplied software, FASTQ files were generated. All reads were aligned to NCBI GRCh37/hg19 using the Burrows-Wheeler alignment (BWA) algorithm v0.7.10. Piccard Tools v1.119 was used for SAM to BAM conversion. The final BAM file was used for variant calling using a custom pipeline MDLVC v.6 (22) and HaplotypeCaller v3.3. Variants with low variant allele frequency (VAF) (<5%) and variants found in a reference pool of normal samples were excluded. Samples (cases 5 and 16) with lower quality sequencing data had an additional VAF filter of 9% applied. Variants meeting quality criteria were then annotated using the COSMIC database v82, dbSNP v150, Annovar (07042018) and Ensembl variant effect predictor(23). Manual review of variant calls was performed using the Broad Institute’s Integrated Genomics Viewer (IGV) v.2.3.4. Tumor mutation burden (TMB) was calculated using a subset of the sequenced genes per published methods (24). Detailed NGS and bioinformatics methods can be found in *Supplementary Information*.

Variant significance was determined by cross-referencing COSMIC (v88), gnomAD (r2.0.2) and ClinVar databases factoring in VAF, functional consequence, level of evidence in the respective databases, evidence of the variant in hematolymphoid malignancies, presence of other variants affecting the same amino acid, and mutational frequency of the gene in hematolymphoid malignancies. Using this rubric, each variant was assigned into one of four categories: likely somatic, cannot exclude somatic/possibly somatic, cannot exclude somatic/possibly germline, and likely germline. Within the “cannot exclude somatic” category, variants were grouped into the possibly germline category if the gene in question had not been reported to be mutated in either dFL (20, 21), pFL (25), cFL (14, 19), or MZL (26). Variants assigned to the “likely somatic” and “cannot exclude somatic/possibly somatic” categories were included for further analyses.

### Tumor Clonality and Cellularity Analysis

Tumor clonality and subclonality analysis was assessed based on several formulas that take into account the admixture of lymphoma cells with normal cells, the presence of clonal and subclonal mutations, and the combined impact of CNVs and coding mutations (see *Supplementary Information*). The most dominant mutation in each tumor, which accounted for the impact of co-occurring CNVs, was used to estimate tumor purity/cellularity. All other variants were divided by this number to derive the normalized subclonal representation of the mutation within the tumor. If tumor purity estimates based on VAF greatly exceeded the morphologic estimate, variants contributing to the over-estimation were re-reviewed for the likelihood of germline derivation and potential for undetected co-occurring CNVs. Should these variants be found, tumor purity was re-calculated accordingly. If this resulted in re-assignment of variants to the possibly/likely germline category, these variants were then excluded from subsequent analyses.

### Statistical Analysis and Graphing

Statistical analyses were performed using GraphPad Prism version 8 (GraphPad Software, La Jolla, CA) and Microsoft Office Excel 2010 (Microsoft, Redmond, WA). Continuous variables were compared using parametric unpaired two-tailed t-tests, while categorical variables were compared using Fisher’s exact test. Detailed statistical analyses are described in the *Supplementary Information*. Mutation representation within protein domains was mapped using MutationMapper (27) and Lollipop (28).

## Results

### dFL shows unique pathologic features

In total, 16 cases of LGFL meeting the inclusion and exclusion criteria were identified. All cases underwent consensus review by the study authors. Summary patient and pathologic findings are detailed in **Table 1**. Histologically, all cases showed ≥75% diffuse growth with many demonstrating microfollicle formation (**Figure 1**), which are miniaturized abnormal follicles predominantly composed of centrocytes lacking follicular dendritic networks. Other notable features include frequent sclerosis and interstitial fibrosis, focal preservation of normal lymph node structures, including normal germinal centers, and more rounded nuclear cytology of the lymphoma cells. Cases with microfollicles tended to have lymphoma cells with centrocyte-like nuclei within microfollicles, and lymphoma cells with more rounded nuclei outside. Immunohistochemical studies confirmed that all lymphomas expressed BCL6, CD10 and CD23, and most expressed variable BCL2. Two cases showed equivocal BCL2 staining (case 7 and 8) due to extensive T-cell admixtures. One case was BCL2 negative (case 15). Some cases also demonstrated disparate staining patterns for BCL2 (cases 6 and 9) and CD10 (case 9) within and outside of microfollicles.

**Figure 1.**
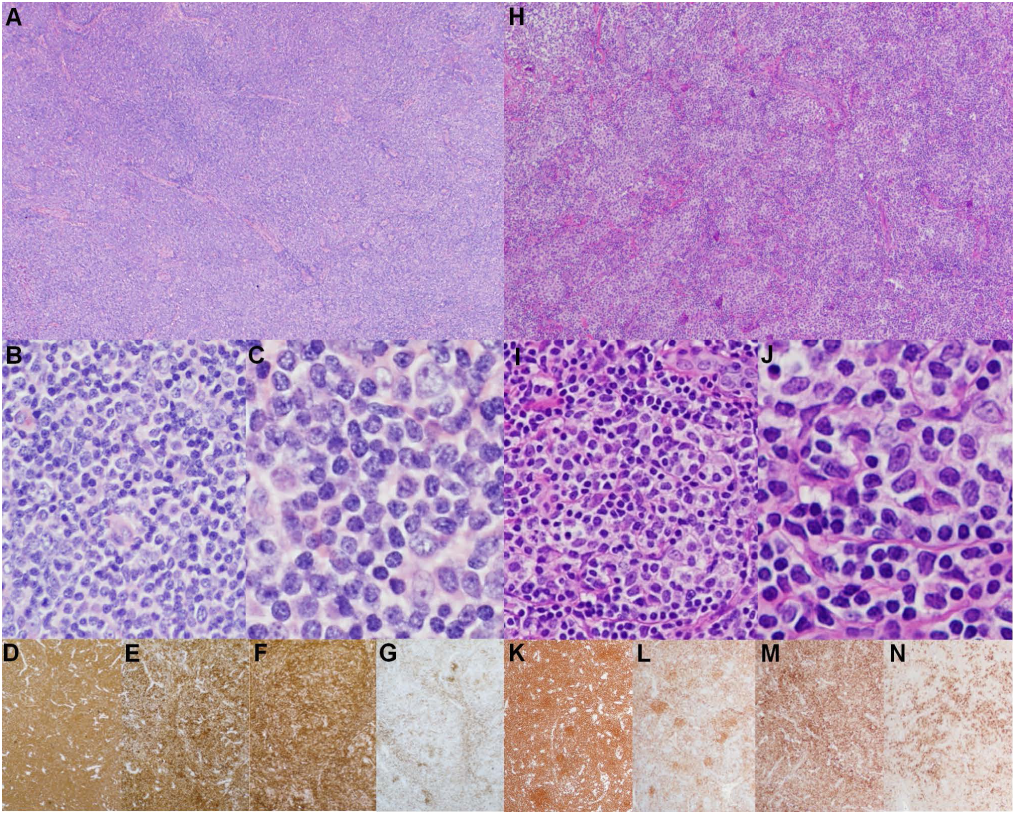
Representative histologic and immunohistochemical features in a case with a purely diffuse growth pattern (Case 1; A-G) and a case with a microfollicular growth pattern (Case 6; H-N). Low power (2X) H&E image demonstrating complete nodal architectural effacement and replacement by a diffuse lymphoid proliferation (**A**) or replacement by a vaguely-nodular proliferation of microfollicles (**H**). 20X H&E image demonstrating that the proliferation comprises a mixture of small lymphocytes and a few scattered large transformed cells (**B, I**). 40X H&E image showing small lymphocytes with rounded nuclear contours (**C**) and small lymphocytes with more angulated and irregular nuclei resembling centrocytes (**J**) that are admixed with occasional large cells resembling centroblasts. Low power (2X) immmunohistochemical images showing staining patterns for CD20 (**D, K**), CD10 (**E, L**), BCL2 (**F, M**), and CD23 (**G, N**).

### Chromosome 16p13 and 1p36 are recurrently altered in the absence of BCL2/IGH

FISH for *IGH/BCL2* was completed for 15 of 16 cases. All interpretable results showed two green (*IGH*) and two orange (*BCL2*) signals without evidence of fusion (**Figure 2C**). SNP-microarray studies were performed on all cases (**Figure 2A** and **2B**) with one case failing quality control. Total CNVs observed per sample ranged from 0-9 (median 2; 95% CI 2-5). Only one sample (case 12) showed no CNVs. Recurrent alterations present in ≥ 4 samples (**Figure 2D**) included loss/CN-LOH of 16p13.3 (9 loss and 1 CN-LOH, 66.7%), loss/CN-LOH of 1p36.3 (4 loss and 3 CN-LOH, 46.7%), gain/CN-LOH of 8q24 (4 gain and 1 CN-LOH, 33.3%), gain of 8p22 (4, 26.7%), and gain/CN-LOH of 8q (3 gains and 1 CN-LOH, 26.7%). Six cases (6/15, 40.0%) showed abnormalities of both 1p36 and 16p13 (**Figure 3**). The minimal deleted region on 16p (16p13.3, 7.1 Mb) contains 238 genes, including *CREBBP*. Nine out of 10 cases with 16p13.3 abnormalities demonstrated slightly larger CNVs that also included *CIITA* and *SOCS1*. The minimal deleted region on 1p (1p36.33-1p36.31, 5.1 Mb) contains 101 genes, including *TNFRSF14*. Full list of CNVs can be found in *Supplemental Table S1*.

**Figure 2.**
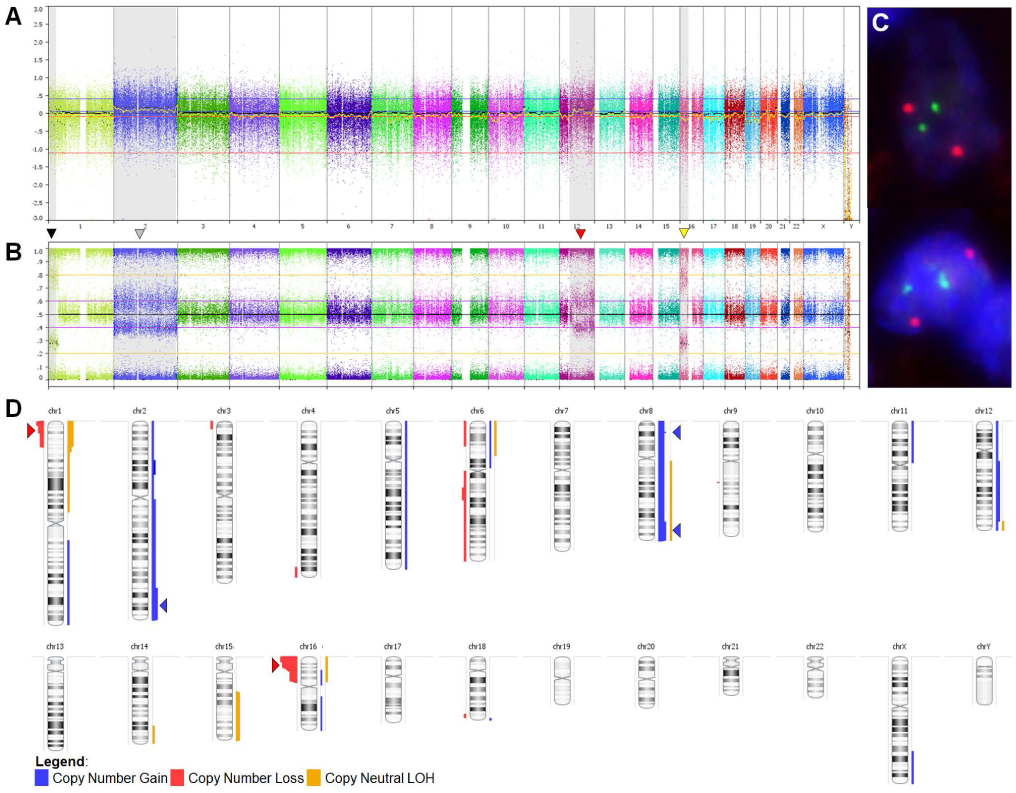
Representative SNP-array (Case 2) and FISH (Case 12) analysis, and chromosomal ideogram showing aggregate copy-number variations. Log-R and smoothed Log-R (gold line) (**A**) and B-Allele frequency plots (**B**) across all chromosomes in a single case. Grey-shaded regions represent observed CNVs, including copy-neutral loss-of-heterozygosity (CN-LOH) of 1p (black arrowhead), gain of chromosome 2 (grey arrowhead), gain of 12q (red arrowhead), and loss of 16p (yellow arrowhead). **C**. FISH analysis using the *IGH/BCL2* dual color fusion probe demonstrating two green and two orange signals, and absence of any fused (yellow) signals that would indicate t(14;18)(q32;q21). **D**. Recurrent CNVs (present in 4+ cases), including loss/CN-LOH of 16p13.3 and loss/CN-LOH of 1p36.3 (red arrowheads), and gain/CN-LOH of 8q24, gain of 8p22 and gain/CN-LOH of 8q (blue arrowheads).

**Figure 3.**
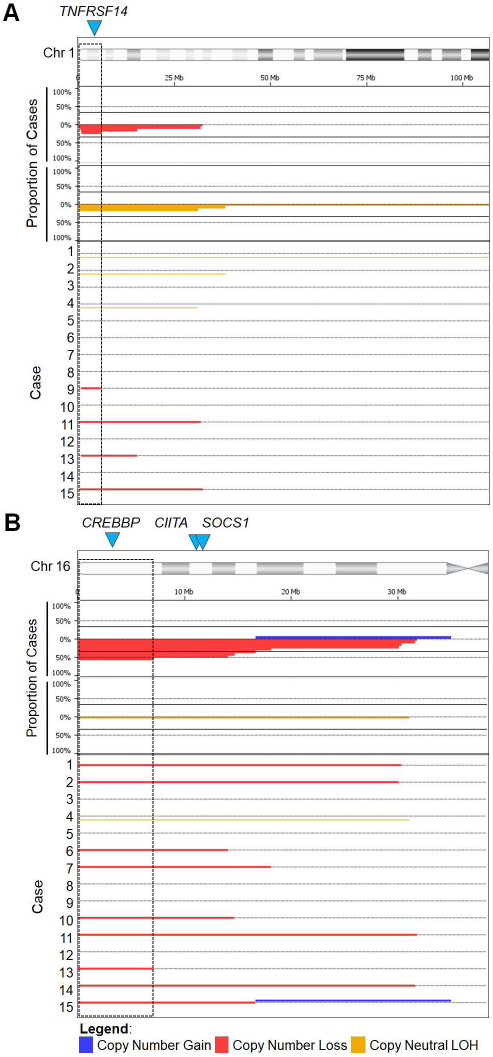
SNP-array profiles of the minimally altered regions on chromosomes 1p (A) and 16p (B). Dashed boxes represent the minimum altered regions of 1p36.33-p36.31 and 16p13.3, respectively. The top panels shows aggregate prevalence of the indicated chromosomal abnormality with horizontal lines representing the aggregate length of the abnormality. The bottom panels demonstrate the CNV observed in each case. Blue arrowheads show the location of overlapping recurrently mutated genes: *TNFRSF14, CREBBP* and *SOCS1*.

### CREBBP and STAT6 are highly recurrently co-mutated

Next-generation sequencing (NGS) was performed in all cases. A total of 161 “likely somatic” and “cannot exclude somatic” variants were identified in 56 genes. 157 (97.5%) of these variants were classified as likely somatic, while 4 (2.5%) were classified as cannot exclude somatic. Clonality and cellularity analysis (see below) reclassified two “likely somatic” variants (*KMT2C* and *SPEN*) and two “cannot exclude somatic” variants (*KMT2C* and *NOTCH1*) as possibly germline, and reclassified the remaining two “cannot exclude somatic” variants (*CSMD3* and *CARD11*) as possibly somatic. Once the possibly germline variants were removed, along with other “likely germline” variants, a total of 157 likely/possibly somatic mutations were identified in 56 genes (**Figure 4**). The number of mutations identified in each case ranged from 6–18 (median 9.5, 95% CI 7–11). Potential aberrant somatic hypermutation (ASHM), suggested by the presence of multiple non-deleterious mutations with similar variant allele frequencies occurring within a single exon and allele, was identified in 3 cases (cases 6, 10 and 11) involving *BCR, SOCS1* and *ACTB* respectively. TMB was calculated for 14 of 16 cases, which showed uniformly low, and occasionally intermediate, TMB ranging from 2.6/Mb to 13.2/Mb (median of 4.4/Mb, 95% CI 2.6-6.1/Mb).

**Figure 4.**
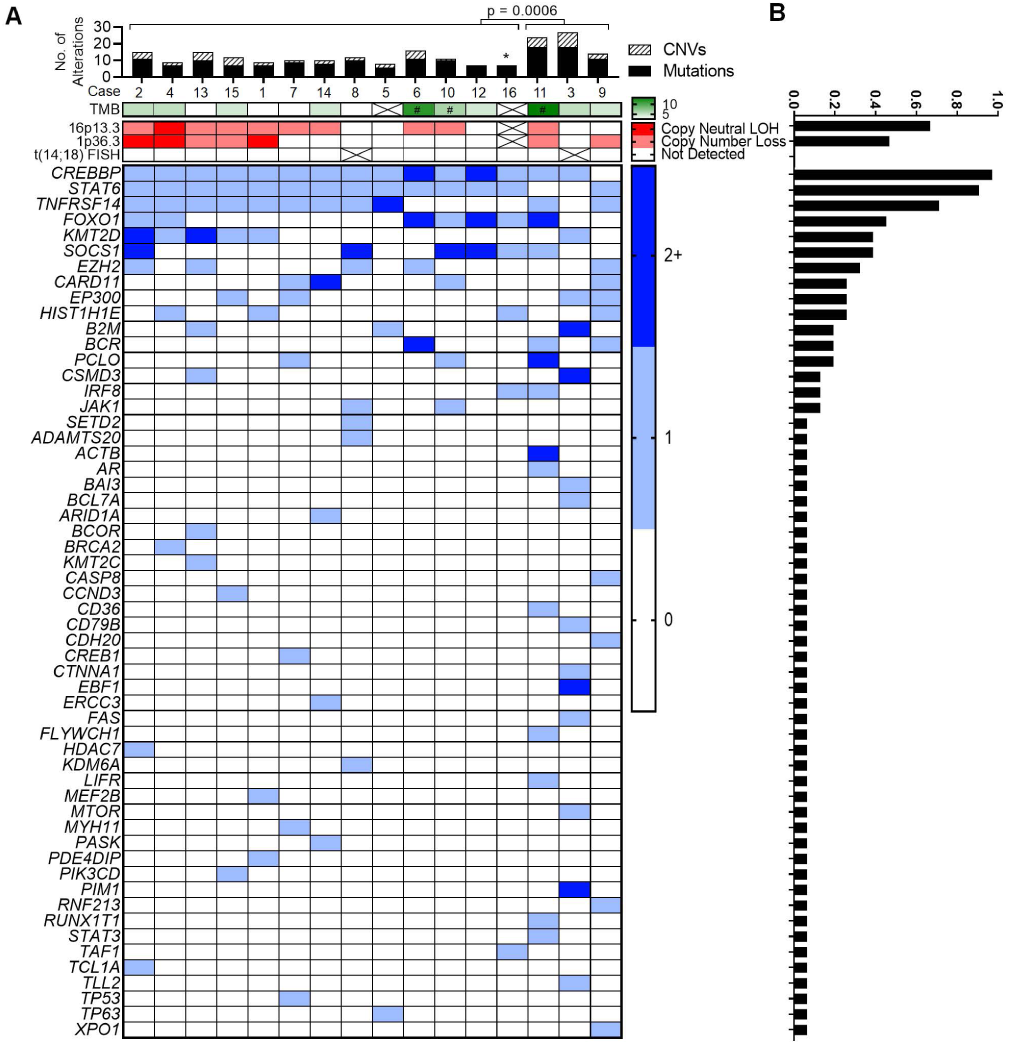
Summary chromosomal and mutational findings. **A**. The top panel shows the total number of mutations (shaded black) and CNVs (shaded with diagonal stripes) detected in each case, and statistical comparison of total alterations per case between the two groups (those with *CREBBP* and *STAT6* co-mutations, and those without)*. The next row in green denotes tumor mutation burden as mutations/Mb. Crossed boxes represent cases without (interpretable) data. The # sign corresponds to cases with suspected aberrant somatic hypermutation. Copy number and FISH abnormalities are summarized in the red panel. Crossed boxes represent cases without (interpretable) data. The bottom panel demonstrates mutational status for all mutated genes. 0 indicates no mutations detected. 1 indicates a single mutation. 2+ indicates 2 or more different mutations within a single case. **B**. Prevalence of the corresponding abnormality in the entire group.*Case 16 was excluded from this analysis due to absence of SNP-array data.

*CREBBP* was nearly-uniformly mutated (15/16 cases, 93.7%) (**Figure 4** and *Supplemental Table S2*). This was followed by *STAT6* (14/16 cases, 87.5%), *TNFRSF14* (11/16 cases, 68.7%), *FOXO1* (11 mutations in 7/16 cases, 43.7%), *KMT2D* (6/16 cases, 37.5%), *SOCS1* (6/16 cases, 37.5%) and *EZH2* (5/16 cases, 31.2%). Incorporating CNV data, 11 of 16 cases (68.7%) showed bi-allelic alterations of 16p13.3 and/or *CREBBP*, and 8 of 16 cases (50.0%) showed bi-allelic alterations of 1p36.3 and/or *TNFRSF14*. Mutations affecting *CREBBP* were mostly missense (12 of 18, 66.7%) or in-frame insertion/deletion (4 of 18, 22.2%) events centered in the HAT histone acetylation protein domain, while mutations affecting *STAT6* were all missense changes occurring in the DNA binding domain (**Figure 5**). Thirteen cases demonstrated *CREBBP* and *STAT6* co-mutation (13/16, 81.2%). Lymphomas carrying mutations in both genes harbored fewer total alterations compared with lymphomas lacking co-mutations (**Figure 4A**). This observation held true for the number of mutations (median 8, ranging from 6–11 vs. median 18, ranging from 12–18; p-value <0.0001), CNVs (median 2, ranging from 0–5 vs. median 6, ranging from 3–9, p-value of 0.0191), and total alterations (median 10, ranging from 7–16 vs. median 24, ranging from 14–27, p-value of 0.0006). Clonality and cellularity assessment (see below) showed that these differences could not be accounted for by lower tumor purity in the co-mutated cases (**Figure 6A**). Even though the number of mutations and CNVs differed between these two groups, TMB did not differ significantly (median 4.4 vs. median 5.3, p-value of 0.1703).

**Figure 5.**
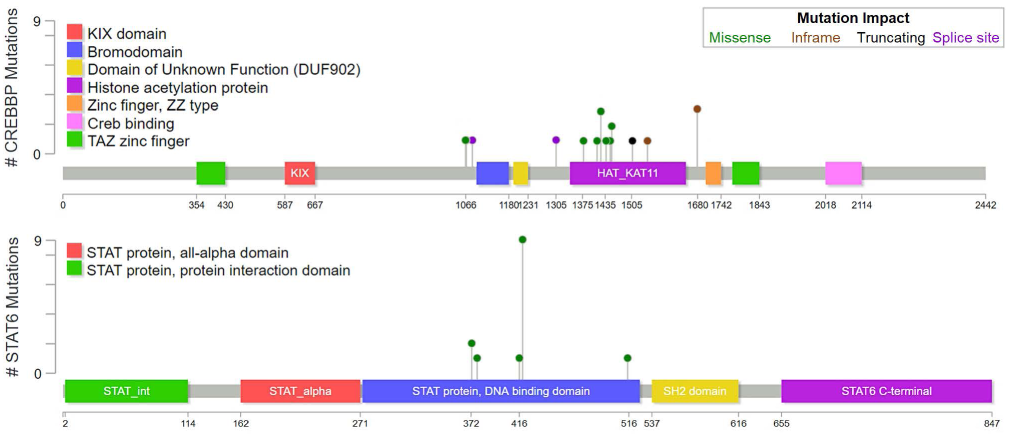
Location and predicted functional impact of mutations in *CREBBP* and *STAT6*. The protein sequence is shown along the x-axis with protein domains marked by colored boxes. The left legend shows the color-coded full name of the domains when full names are abbreviated in the protein depiction. Each circle corresponds to a specific mutation with the height of the circle on the y-axis representing the number of occurrences of that mutation in the studied cases. The color of the dot corresponds to the predicted functional impact of the mutation (top right box).

**Figure 6.**
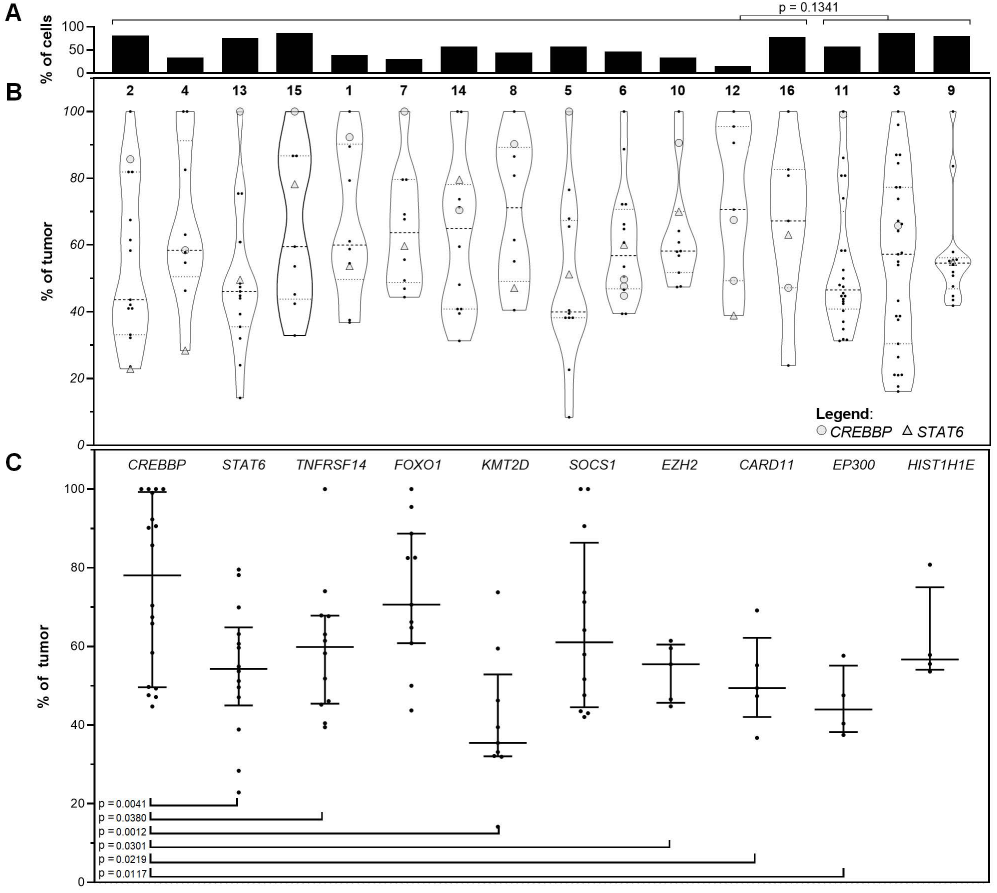
Clonality analysis based on CNVs and mutations. **A**. Tumor purity/cellularity, as calculated by the most dominant CNV/mutation. Cases with *CREBBP* and *STAT6* co-mutations show no significant difference in tumor purity when compared to cases without. **B**. Clonal architecture of individual CNVs and mutations represented as proportion of the tumor cells (normalized to tumor %) on the y-axis. *CREBBP* and *STAT6* mutations are indicated as shaded circles and triangles, respectively. Other mutations are represented as black dots **C**. Individual mutations found in the top ten recurrently mutated genes shown as a proportion of the respective tumor % on the y-axis. Error bars represent median and interquartile range. Statistical analysis of clonal dominance show statistically significant differences between clonal dominance of *CREBBP* versus subclonality of *STAT6, TNFRSF14, KMT2D, EZH2*, and *CARD11* and *EP300*.

### CREBBP mutations are clonally dominant

Integrating both mutation VAFs and (co-occurring) CNV B-allele frequencies, the cellular representation of individual alterations was calculated (**Figure 6B**), which enabled estimation of tumor purity/cellularity (**Figure 6A**), and more accurate variant significance classification. This analysis showed that *CREBBP* mutations are dominant clonal events in most cases (**Figure 6C**) accounting for 78.1% of tumor cells (median 95 CI 49.7-99.0%). In contrast, other recurrently mutated genes frequently represented subclonal events with *STAT6* mutations accounting for 54.3% of tumor cells (median 95% CI 38.9-70.0%, p = 0.0041), *TNFRSF14* mutations accounting for 59.9% of tumor cells (median 95% CI 45.2-67.9%, p = 0.0380), *KMT2D* mutations accounting for 35.5% of tumor cells (median 95% CI 32.0-59.5%, p = 0.0012), *EZH2* mutations accounting for 55.5% of tumor cells (median 94% CI 44.8-61.5%, p = 0.0301), *CARD11* mutations accounting for 49.4% of tumor cells (median 94% CI 36.8-69.2%, p = 0.0219), and *EP300* mutations account for 44.0% of tumor cells (median 88% CI 37.5-57.6%, p = 0.0117).

### CREBBP and STAT6 co-mutation and 16p13 and 1p36 loss represent unique features of dFL

In order to determine if the recurrent CNV and mutational findings from the present study are enriched in dFL, a detailed literature review was performed (**Table 2)**. Each recurrent, and select combinations of, alteration found in the current report was pooled with less comprehensive analyses from 3 previous studies of dFL (3, 20, 21) to identify unique features of dFL. Of note, 2 cases from the Siddiqi *et al* study were not included, as those cases had demonstrable *BCL2/IGH* rearrangements. The aggregate frequencies of particular alterations found in dFL were contrasted with previously published reports for cFL (4-19) and MZL (26, 29-41). Although the previous studies describing CNVs in cFL used a variety of techniques, most of these studies (8/13, 61%) were performed using SNP-array platforms similar to the present method indicating that the results obtained in these prior studies should be comparable to our findings. Compared to cFL and MZL, 16p13 and/or 1p36 abnormalities are far more frequent in dFL. *CREBBP* mutations are slightly more common in dFL, and *STAT6* mutations are much more common in dFL. *CREBBP* and *STAT6* co-mutation is particularly enriched in dFL. All recurrent alterations found in dFL are statistically significantly under-represented in MZL (**Table 2**).

## Discussion

The present study of 16 cases of dFL is the largest series to include detailed pathologic, chromosomal and NGS analyses that reveal novel, and unifying, pathologic and genetic findings. Not only do our findings support continued classification of dFL a variant of cFL, our findings also show how comprehensive molecular profiling can aid in the differential diagnosis and workup of low-grade B-cell lymphoma (LGBCL).

Pathologic analyses identified novel morphologic features of dFL, such as frequent sclerosis, microfollicle formation and rounded nuclear cytology, in addition to the known features of diffuse histology, focal preservation of normal lymph node structures, co-expression of CD23, and variable expression of BCL2 (3, 20). Microfollicles lack follicular dendritic networks rendering them distinct from typical follicles/nodules found in cFL. To our knowledge, this growth pattern has only been associated with dFL (1), and has not been described in any other type of LGBCL to date. Although CD23 was used as a selection criteria for 6 of 16 cases, all cases showed CD23 co-expression suggesting that this may be a unifying feature of dFL, whereas cFL is only occasionally CD23 positive (42, 43). Variable BCL2 expression in the absence of *BCL2/IGH* rearrangements suggests alternative mechanisms of BCL2 up-regulation on the DNA (44, 45) or transcriptional (46, 47) level, although we did not find either copy number gains of the BCL2 locus or mutations of BCL2 in dFL.

Our data demonstrated new associations of loss/CN-LOH of 16p13 and CN-LOH of 1p36, and confirmed the reported absence of t(14;18) and recurrent loss of 1p36 (3, 20). However, the frequency 1p36 abnormalities in our series was far lower than originally published (3), but is similar to the rate reported by Siddiqi *et al* (20). This difference may be related to selection, sampling, and/or technical biases. Unlike the Katzenberger *et al* (3) report where CD23 positivity was found in approximately two-thirds of the lymphomas, there was uniform expression of CD23 in this and the Siddiqi *et al* series, which could skew the distribution of the genetic findings. Alternatively, with larger numbers of dFL being studied, the full spectrum of chromosomal abnormalities is emerging unmasking a lower prevalence of 1p36 deletion. Finally, technical bias could also account for these differences, as the original report used FISH, which has a superior analytical sensitivity (5% of nuclei) to both aCGH and SNP-microarray. While all of the cases we studied had at least 15% tumor cells, it is plausible that subclonal loss of 1p36 may be missed by our approach. Irrespective of the reason for this discordance, combined data suggest that loss of 1p36 alone is not sufficient, or specific, for dFL, especially if array-based techniques or NGS are used. Unlike previous studies, the most predominant CNV observed in our series was loss/CN-LOH of 16p13, which was only found in 2 cases (22.2%) in the Siddiqi *et al* study (20). This apparent discrepancy may, again, be technique related, as array CGH (aCGH) used by Siddiqi *et al* typically shows inferior analytical sensitivity, and cannot detect CN-LOH. Not only are 16p13 abnormalities a novel association in dFL, we also found that the minimally altered region(s) encompassed *CREBBP, CIITA* and *SOCS1*, which suggests a possible cooperative mechanism for tumor immune evasion (19, 48).

Targeted NGS showed near-uniform mutations of *CREBBP* and *STAT6* with clonal dominance of the *CREBBP* mutations suggestive of a founder event. The mutational profiles of dFL in our series showed frequent mutations in genes implicated in GCB derived lymphomas (11, 12), including *CREBBP, TNFRSF14, KMT2D* and *EZH2*, which offers genetic confirmation for the current classification of dFL as a FL variant. We did not identify MAPK pathway mutations associated with pFL (25, 49) indicating dFL shares more genetic similarities with cFL than pFL. Unlike cFL, where t(14;18) represents the founder event (13) and *CREBBP* mutations represent subsequent driver events, our data suggest *CREBBP* mutations represent founder events in dFL in the absence of *BCL2/IGH* rearrangements.

With regard to *CREBBP* mutations, the enrichment for non-truncating mutations within the HAT domain, which leads to enzymatic loss of protein function (12), is similar to what has been previously described in cFL (50). Unlike previous reports of cFL or GCB DLBCL (12, 14), which show majority mono-allelic loss of *CREBBP*, our series identified majority bi-allelic loss of *CREBBP*. In mice, heterozygous/haploinsufficient loss of *CREBBP* coupled with BCL2 over-expression in B-cells leads to the development of GCB lymphomas (50). Without *BCL2/IGH*, however, other anti-apoptotic mechanisms may be implicated in dFL, such as *STAT6* co-mutation. *STAT6* is commonly mutated in classical Hodgkin lymphoma (32%) (51) and PMBL (36%) (52), but is not typically mutated in GCB lymphomas (11-19). Our data show that *STAT6* mutations are always co-mutated with *CREBBP*, or *EP300* that forms the CREBBP/EP300 complex, in dFL, and are frequently associated with concurrent loss of *SOCS1*. The conspicuous co-occurrence of these alterations suggest a degree of cooperativity. Similar to previous studies of GCB lymphoma (53, 54), all of the detected *STAT6* mutations in dFL were missense changes occurring in the DNA binding domain, which has been shown to activate JAK/STAT signaling (53, 54). An important STAT6 target is the *BCL-xL/BCL2L1* (BCL2-like anti-apoptotic protein) gene (55), which is often amplified in epithelial malignancies (56). In PMBL, over-expression pSTAT6 leads to accumulation of BCL-xL (57), a phenomenon that may be reversed by inducing the STAT6 negative regulator SOCS1 (51, 57). The concurrent gain of function of STAT6 and loss of its negative regulator, SOCS1, in dFL may drive high levels of BCL-xL that could serve as a functional surrogate for BCL2 excess to cooperate with *CREBBP* bi-allelic loss in the development of dFL. Future studies could evaluate the possibility that *CREBBP* loss and *STAT6* gain, possibly through BCL-xL, are sufficient to induce dFL-like lymphomas.

A major limitation of this study is that it is correlative, and lacks functional confirmation of the findings and the proposed interactions. Another limitation is the small sample size and lack of clinical follow-up, which is a consequence of the exceedingly rare occurrence of this lymphoma, and the frequent extramural consultative nature of the pathology review. As described earlier, the uniform inguinal location and CD23 positivity found in the present study may bias the results towards apparent unifying pathologic and molecular features. Given that these two features are commonly used as criteria to diagnose this variant of FL, only a large-scale screen of diffuse-pattern LGFL would allow identification of sufficient numbers of CD23-negative/non-inguinal cases to investigate this possibility. Although some t(14;18)-negative FL have *BCL6* abnormalities (translocations or amplification), we did not pursue *BCL6* translocations since that was not a criterion used in the original Katzenberger *et al* (3) definition, and we did not find *BCL6* amplification in our series. Additional limitations are technical in nature. There may be false negativity, in particular for subclonal 1p36 deletion, due to low tumor cellularity seen in a small number of cases. The lack of matched germline tissue can confound tumor-only SNP-microarray and NGS analysis, although we have detailed conservative and comprehensive interpretive guidelines to limit misattribution of germline variants as somatic mutations. Lastly, we did not perform detailed genetic analyses of a control group comprising cFL and MZL to determine if the CNVs and mutations found in dFL are truly enriched by a direct case-control comparison. Since a broad range of techniques and analysis methods were used by the referenced studies, there may be apparent differences in chromosomal and mutational patterns that is simply methodology-related. However, since many of the referenced studies used very similar techniques to the ones used in the present study, and reproducible molecular patterns were identified through this review, the presented aggregate re-analysis of the literature should represent a reliable estimate of the true rates of chromosomal and molecular abnormalities found in cFL and MZL, from which dFL differ.

Combined with the previously published studies, 66 dFL cases have now been pathologically and genetically characterized. As the WHO classification moves towards molecularly-defined lymphoma entities, such as pFL, the unifying pathologic and genetic features described herein may aid in the accurate subclassification of LGFL. The diagnostic distinction between the dFL from cFL with prominent diffuse growth is specifically recommended by the 2016 WHO (1) when an excisional biopsy is available, as the former will consistently lack t(14;18) *BCL2/IGH* rearrangements. In diagnostically challenging cases, the ancillary work-up should begin with FISH. Once absence of *BCL2* rearrangement is confirmed, NGS and CNV detection should follow. Identification of the characteristic 1p36 and/or 16p13 abnormalities along with *CREBBP* and *STAT6* co-mutations would support a diagnosis of a t(14;18)-negative dFL. The present literature, including our findings, has identified genetically distinct profiles of subtypes of LGBCL, which support the incorporation of genomic studies in the routine lymphoma workup, as the field moves towards molecular classification of lymphoma subtypes.

## Acknowledgements

The authors would like to thank the patients, and submitting pathologists and clinicians who provided supporting case data and patient samples for the study. This work was supported in part by the Sidney Kimmel Comprehensive Cancer Center Grant from the National Institutes of Health (P30 CA006973; R.R.X., A.S.D., C.D.G.) and by the intramural research program of the Center for Cancer Research, National Cancer Institute (Y.X., S.P., E.S.J.). We would also like to thank the University of California, Los Angeles, Clinical Cytogenetics Laboratory and Dr. Nagesh Rao for providing usage of the Nexus Copy Number chromosomal analysis software.

## Authorship Contributions

R.R.X and C.D.G. designed the study, analyzed the data, and wrote the manuscript; L.M.H and R.Y. performed the SNP-array and NGS studies, and FISH studies respectively, contributed to the manuscript, and critically reviewed the manuscript. A.P. developed the NGS data analysis and TMB pipelines, assisted with data analysis, contributed to the manuscript, and critically reviewed the manuscript. Y.X., S.P. and E.S.J. contributed to the study design, analyzed the data, and critically reviewed the manuscript. A.S.D., C.M.M. and S.M.F.G. contributed to the study design, reviewed the data, and critically reviewed the manuscript.

---

**Table 1. Patient demographic and pathologic characteristics**.

**Table 2. Recurrently detected copy number variants and mutations found in diffuse follicular lymphoma (dFL) compared to previously published studies of conventional follicular lymphoma (cFL) and marginal zone lymphoma (MZL).** CNVs in dFL detected in the present report and previously published studies* (3, 20) were grouped and compared to previously published studies of cFL (4-10) and MZL (26, 29, 38-41). Mutations in dFL detected in the present report and previously published studies* (20, 21) were grouped and compared to mutations found in previously published studies of cFL (11-19) and MZL (26, 29-37). The prevalence of each alteration in dFL, cFL and MZL are shown along with the total number of samples studied. Statistical significance for each alteration found in dFL is tested against the same rates in cFL and MZL, and the resultant p-values are shown.

*Two cases were removed from a previously published study (20) of dFL in this aggregate analysis due to the presence of t(14;18) in those cases.

