## Supplementary material for "*CREBBP* and *STAT6* co-mutation and 16p13 and 1p36 loss define the t(14;18)-negative diffuse variant of follicular lymphoma": Table 2

**Table 2. Recurrently detected copy number variants and mutations found in diffuse follicular lymphoma (dFL) compared to previously published studies of conventional follicular lymphoma (cFL) and marginal zone lymphoma (MZL).**

| **Copy Number Loss/CN-LOH** | **dFL % (N)** | **cFL % (N)** | **p-value** | **MZL % (N)** | **p-value** |
| --- | --- | --- | --- | --- | --- |
| 1p36 | 76.4% (42/55) | 21.8% (158/724) | <0.0001 | 6.5% (47/726) | <0.0001 |
| 16p13 | 38.5% (10/26) | 9.1% (53/583) | <0.0001 | 7.6% (30/396) | <0.0001 |
| 1p36+16p13 | 30.8% (8/26) | 3.4% (8/233) | <0.0001 | 2.2% (4/178) | <0.0001 |
| **Mutation** | **dFL % (N)** | **cFL % (N)** | **p-value** | **MZL % (N)** | **p-value** |
| *CREBBP* | 87.1% (27/31) | 59.6% (374/627) | 0.0021 | 6.9% (37/534) | <0.0001 |
| *STAT6* | 87.1% (27/31) | 10.4% (61/528) | <0.0001 | 0.6% (2/309) | <0.0001 |
| *TNFRSF14* | 54.8% (17/31) | 31.1% (202/649) | 0.0094 | 3.1% (10/327) | <0.0001 |
| *FOXO1* | 29.0% (9/31) | 7.4% (30/331) | 0.0027 | 1.3% (4/309) | <0.0001 |
| *KMT2D* | 48.4% (15/31) | 76.9% (464/603) | 0.0009 | 11.2% (67/597) | <0.0001 |
| *SOCS1* | 19.4% (6/31) | 3.0% (12/270) | 0.0056 | 1.0% (3/309) | <0.0001 |
| *EZH2* | 29.0% (9/31) | 19.2% (123/641) | 0.1716 | 1.5% (7/469) | <0.0001 |
| *CREBBP+STAT6* | 74.2% (23/31) | 7.4% (35/421) | <0.0001 | 0.6% (2/309) | <0.0001 |
| *CREBBP+STAT6+TNFRSF14* | 38.7% (12/31) | 2.8% (13/421) | <0.0001 | 0.3% (1/309) | <0.0001 |
| *CREBBP/EP300+KMT2D* | 41.9% (13/31) | 56.0% (314/561) | 0.1402 | 2.2% (11/502) | <0.0001 |
