## Supplementary Information for "*CREBBP* and *STAT6* co-mutation and 16p13 and 1p36 loss define the t(14;18)-negative diffuse variant of follicular lymphoma"

**Supplemental Methods**

**Pathologic Case Selection**

A formal sample size analysis was not performed since this is a case series study of an extremely rare entity, for which little is known regarding associated chromosomal and molecular abnormalities. There was no pre-conceived effect size being predicted, and no power estimates were performed. After appropriate institutional review board approval at both sites (Johns Hopkins Hospital and the National Cancer Institute) cases with available formalin-fixed paraffin embedded (FFPE) tissue were selected. Informed consent was not feasible, required, or obtained due to the retrospective nature of the study. Photomicrographs were taken using an Olympus BX46 microscope with an Olympus DP72 camera, or an Olympus BX50 microscope with an Olympus DP71 camera (Olympus Lifescience, Tokyo, Japan). All images were acquired using Cellsens (Olympus Lifescience, Tokyo, Japan), and modified in Adobe Photoshop (Adobe, San Jose, CA).

**Fluorescence In-Situ Hybridization**

Prior to probe hybridization, the unstained slides were deparaffinized following pretreatment with Protease I on the VP 2000 processor (02J08-032, Abbott Molecular, Des Plaines, IL). After this, the slides and the probe (Vysis LSI IGH/BCL2 dual color dual fusion probe, Abbott Molecular, Des Plaines, IL, Catalogue #05J71-001) were co-denatured at 75 °C for 7 minutes and allowed to anneal for 22 hours at 37 °C in a humidified atmosphere. At the end of this incubation the slides were washed in 2X SSC/0.3% NP-40 for 2 minutes at 72 °C, then 2 minutes at room temperature, and a final wash using 2X SSC at room temperature. The slides were then counterstained with DAPI and a cover slip was applied using VECTASHIELD mounting medium (H-1000, Vector Laboratories, Burlingame, CA). A fluorescence microscope was used to evaluate probe patterns on each slide for the presence or absence of fusion signals. Since the probes comprise a mixture of two differentially labeled probes designed to specifically detect the t(14;18)(q32;q21) translocation, a fusion would result in a yellow signal indicating juxtaposition of the orange fluorochrome direct-labeled *BCL2* locus next to the green fluorochrome direct-labeled *IGH* locus. Cells containing 2 green and 2 orange signals would indicate absence of an *IGH/BCL2* translocation event. FISH using this probe was performed in singlicate for each sample.

**SNP-Microarray Analysis Interpretation**

CNVs larger than 3 megabases (Mb) with a corresponding B-allele frequency of at least 0.45 or less, or 0.55 or greater, were considered positive findings. Smaller changes, less than 3 Mb, were rarely annotated as present when B-allele frequencies deviated greatly from 0.5. CNVs with positive smoothed log-R were designated as gain, negative smoothed log-R were designated as losses, and neutral smoothed log-R were designated as copy-neutral loss-of-heterozygosity (CN-LOH), also known as acquired uniparental disomy (aUPD) in the somatic setting. This analysis allowed for positive identification of lymphoma-related CNVs when lymphoma cells comprise approximately 20% of cells or greater. CN-LOH events with very high BAF likely representing germline constitutional changes were not included in subsequent analyses. While we cannot completely exclude the presence of germline clonal mosaicism, the population frequency of these events is exceedingly low (0.5-1.5%) (1), and we do not believe we would have captured these rare events in a cohort of 16 patients. SNP-microarray was performed in singlicate for each sample.

**Next-Generation Sequencing**

NGS for this study was performed using a laboratory-developed test that was previously clinically validated by the CLIA-certified Molecular Diagnostics Laboratories at Johns Hopkins in concordance with standards recommend by the College of American Pathologists. NGS was performed in singlicate for each sample. In addition to determining DNA quality/quantity input requirements, assay-specific artifacts (“black-list”), NGS data acceptance criteria, and bioinformatics pipeline performance, these studies also performed orthogonal variant confirmation by Sanger sequencing and/or pyrosequencing. The validated limit of detection for single nucleotide variants was >=2.0% variant allele frequency (VAF) and for insertion/deletions was >= 3.0% VAF. Additional assay characteristics are as follows.

| **Variant Type** | **Sensitivity** | **Specificity** |
| --- | --- | --- |
| Single nucleotide substitutions | 98% | >99% |
| Insertion/deletion (1-40 bp) | >95% | >99% |

| **Reproducibility** | **Type** | **Value** |
| --- | --- | --- |
| Concordance between replicates | Inter-run | 99% |
| Concordance between replicates | Intra-run | 99% |

Briefly, sequencing libraries were prepared using the Agilent SureSelect‐XT Target Enrichment Kit. Between 0.2 and 1 μg of DNA was fragmented to a size of 250 to 300 bp using a Covaris M220 sonicator (Covaris, Inc, Woburn, MA). The DNA fragments were end‐repaired and A‐tailed, then adaptors were added by ligation and the fragments were enriched by PCR (six cycles). Each library was then hybridized to a SureSelect custom panel 2.8 M bait set (Agilent) according to the manufacturer's protocol. After stringent washing, the captured DNA was amplified with 12 cycles of PCR per manufacturer's protocol. The size and concentration of captured DNA was assessed using a Tapestation 2200 (Agilent). All captured samples were clustered on a cBOT system, and sequenced on a single lane of a PE‐flow cell on a HiSeq2500 (Illumina), using a 2 × 150 bp PE protocol. The following HiSeq operating software versions were used: HiSeq Control Software HCS 2.2.68, Real Time Analysis Software RTA 1.18.66, HiSeq Recipe Fragments v1.5.21, .NET 4.5.1, Sequencing Analysis Viewer-SAV v1.8.46, and Batch Installer v1.4. FASTQ formatted raw data files were generated by the vendor supplied bcl2fastq v1.8.4 software (Illumina Biotechnology, San Diego, CA).

After FASTQ file generation, all reads were aligned to the NCBI genome build 37 (GRCh37.p13/hg19) using the Burrows-Wheeler alignment (BWA) algorithm v0.7.10. Piccard Tools v1.119 was used for SAM to BAM conversion. The final Binary Alignment Map (BAM) file was used for variant calling using a custom variant caller pipeline MDLVC v.6 and an additional variant caller HaplotypeCaller v3.3, which called variants and applied multiple exclusionary filters (≤4 variant reads in both directions, total read depth ≤50x, variant allele frequency ≤5%, common germline single-nucleotide polymorphisms based on dbSNP, strand bias, Qscore ≤10, and known laboratory validated artifacts). Only variants called by both variant callers were considered true variants. Additionally, all variants were filtered against a pool of normal reference samples (N=16) using the following criteria: any variants with a variant allele frequency (VAF) within 3 standard deviations from the mean VAF seen in the pool of normal reference samples are filtered out. Manual review was performed using IGV (v2.3.4), and included hot-spot regions of *CREBBP* and *STAT6*.

Tumor mutation burden (TMB) was calculated using a subset of the sequenced genes (N=432) that totaled 1.14 MB of sequence per previously published methods (2), which was derived by analyzing 211 internal tumor NGS results against the gold-standard method. These studies demonstrated a correlation coefficient of 0.91 between our method and the gold-standard method for TMB. Briefly, final variant calls that passed the above filters were additionally filtered to calculate TMB. Variants were excluded based on the presence in the ExAC v0.3.1 database, presence as a “likely somatic” change in the COSMIC v82 database, location in non-coding regions (intronic, 5’UTR, 3’UTR, intronic splice variants, non-coding RNA sequences), and designation as “likely germline” based on the SGZ algorithm (3), a computational method that takes into account estimated tumor purity to designate likely germline, ambiguous, and likely somatic origin of a variant. Tumor purity for the SGZ algorithm was estimated based on CNVs, or based on morphology when no CNVs were detected. TMB was calculated as the number of remaining variants per Mb of sequence evaluated. Using this method, previous analysis of 916 tumors of various organs, showed that TMB of ≤5.28/Mb would be considered low, TMB of ≥20.22/Mb would be considered high, and TMB of 5.29-20.21/Mb would be considered intermediate.

**Genes Assessed by Next-Generation Sequencing**

The 641 gene sequencing panel includes: *ABL1, ABL2, ACTB, ACVR1B, ACVR2A, ADAMTS20, AFF1, AFF3, AKAP9, AKT1, AKT2, AKT3, ALK, APC, APH1A, AR, ARAF, ARFRP1, ARHGAP26, ARID1A, ARID1B, ARID2, ARNT, ASMTL, ASMTL, ASXL1, ATF1, ATM, ATR, ATRX, AURKA, AURKB, AURKC, AXIN1, AXL, B2M, BAI3, BANF1, BAP1, BARD1, BCL10, BCL11A, BCL11B, BCL2, BCL2L1, BCL2L2, BCL3, BCL6, BCL7A, BCL9, BCOR, BCORL1, BCR, BIRC2, BIRC3, BIRC5, BLM, BLNK, BMPR1A, BRAF, BRCA1, BRCA2, BRD3, BRD4, BRIP1, BRSK1, BTG2, BTK, BTLA, BUB1B, C11ORF30, CAD, CALR, CARD11, CASC5, CASP8, CBFB, CBL, CBLB, CCND1, CCND2, CCND3, CCNE1, CCT6B, CD22, CD274, CD36, CD58, CD70, CD79A, CD79B, CDC73, CDH1, CDH11, CDH2, CDH20, CDH5, CDK12, CDK4, CDK6, CDK8, CDKN1B, CDKN2A, CDKN2B, CDKN2C, CEBPA, CEBPB, CHD2, CHEK1, CHEK2, CHIC2, CIC, CIITA, CKS1B, CMPK1, COL1A1, CPS1, CRBN, CREB1, CREBBP, CRKL, CRLF2, CRLF2, CRTC1, CSF1R, CSF3R, CSMD3, CTCF, CTNNA1, CTNNB1, CUX1, CXCR4, CYLD, CYP2C19, CYP2D6, DAXX, DCC, DDB2, DDIT3, DDR2, DDX10, DDX3X, DEK, DICER1, DNM2, DNMT1, DNMT3A, DOT1L, DPYD, DST, DTX1, DUSP2, DUSP9, EBF1, ECT2L, EED, EGFR, ELP2, EML4, EP300, EP400, EPHA3, EPHA5, EPHA7, EPHB1, EPHB4, EPHB6, ERBB2, ERBB3, ERBB4, ERCC1, ERCC2, ERCC3, ERCC4, ERCC5, ERG, ESR1, ETNK1, ETS1, ETV1, ETV4, ETV6, EXOSC6, EXT1, EXT2, EZH2, FAF1, FAM123B, FAM46C, FANCA, FANCC, FANCD2, FANCE, FANCF, FANCG, FANCL, FAS, FBXO11, FBXO31, FBXW7, FGF10, FGF14, FGF19, FGF23, FGF3, FGF4, FGF6, FGFR1, FGFR2, FGFR3, FGFR4, FH, FHIT, FLCN, FLI1, FLT1, FLT3, FLT4, FLYWCH1, FN1, FOXL2, FOXO1, FOXO3, FOXP1, FOXP4, FRS2, FUBP1, FZR1, G6PD, GADD45B, GATA1, GATA2, GATA3, GDNF, GID4, GNA11, GNA12, GNA13, GNAQ, GNAS, GPR124, GRIN2A, GRM8, GSK3B, GSX2, GTSE1, GUCY1A2, H3F3A, HCAR1, HDAC1, HDAC4, HDAC7, HGF, HIF1A, HIST1H1C, HIST1H1D, HIST1H1E, HIST1H2AC, HIST1H2AG, HIST1H2AL, HIST1H2AM, HIST1H2BC, HIST1H2BJ, HIST1H2BK, HIST1H2BO, HIST1H3B, HLF, HNF1A, HOOK3, HOXA11, HOXA13, HOXA9, HRAS, HSP90AA1, HSP90AB1, ICK, ID3, IDH1, IDH2, IGF1R, IGF2, IGF2R, IKBKB, IKBKE, IKZF1, IKZF2, IKZF3, IL2, IL21R, IL6ST, IL7R, ING4, INHBA, INPP4B, INPP5D, IRF1, IRF4, IRF8, IRS2, ITGA10, ITGA9, ITGB2, ITGB3, JAK1, JAK2, JAK3, JARID2, JUN, KAT6A, KAT6B, KDM2B, KDM4C, KDM5A, KDM5C, KDM6A, KDR, KEAP1, KIT, KLF4, KLF6, KLHL6, KMT2A, KMT2B, KMT2C, KRAS, LAMP1, LCK, LEF1, LIFR, LMO1, LPHN3, LPP, LRP1B, LRRK2, LTF, LTK, MAF, MAFB, MAGEA1, MAGED1, MAGI1, MALT1, MAML2, MAP2K1, MAP2K2, MAP2K4, MAP3K1, MAP3K14, MAP3K6, MAP3K7, MAPK1, MAPK8, MARK1, MARK4, MBD1, MCL1, MDM2, MDM4, MED12, MEF2B, MEF2C, MEN1, MET, MIB1, MITF, MKI67, MKL1, MLH1, MLL, MLL2(KMT2D), MLL3, MLLT10, MLLT4, MLLT6, MMP2, MN1, MNX1, MPL, MRE11A, MSH2, MSH3, MSH6, MTOR, MTR, MTRR, MUC1, MUTYH, MYB, MYC, MYCL1, MYCN, MYD88, MYH11, MYH9, MYO18A, MYST3, NBN, NCOA1, NCOA2, NCOA3, NCOA4, NCOR1, NCOR2, NCSTN, NF1, NF2, NFE2L2, NFKB1, NFKB2, NFKBIA, NIN, NKX2-1, NLRP1, NOD1, NOTCH1, NOTCH2, NOTCH4, NPM1, NRAS, NSD1, NT5C2, NTRK1, NTRK2, NTRK3, NUMA1, NUP214, NUP93, NUP98, P2RY8, P2RY8, PAG1, PAK3, PALB2, PARP1, PASK, PAX3, PAX5, PAX7, PAX8, PBRM1, PBX1, PC, PCBP1, PCLO, PDCD1, PDCD11, PDCD1LG2, PDE4DIP, PDGFB, PDGFRA, PDGFRB, PDK1, PER1, PGAP3, PHF6, PHOX2B, PICALM, PIGA, PIK3C2B, PIK3CA, PIK3CB, PIK3CD, PIK3CG, PIK3R1, PIK3R2, PIM1, PKHD1, PLAG1, PLCG1, PLCG2, PLEKHG5, PML, PMS1, PMS2, POT1, POU5F1, PPARG, PPP2R1A, PRDM1, PRDM16, PRKAR1A, PRKDC, PRSS8, PSIP1, PTCH1, PTEN, PTGS2, PTPN11, PTPN2, PTPN6, PTPRD, PTPRO, PTPRT, RAD21, RAD50, RAD51, RAF1, RALGDS, RARA, RASGEF1A, RB1, RBM15, RECQL4, REL, RELN, RET, RHOA, RHOH, RICTOR, RIT1, RNASEL, RNF2, RNF213, RNF43, ROS1, RPN1, RPS6KA2, RPTOR, RRM1, RUNX1, RUNX1T1, S1PR2, SAMD9, SBDS, SDHA, SDHB, SDHC, SDHD, SEPT9, SERP2, SETBP1, SETD2, SF3B1, SGK1, SH2B3, SH2D1A, SKP2, SMAD2, SMAD4, SMARCA1, SMARCA4, SMARCB1, SMC1A, SMC3, SMO, SMUG1, SOCS1, SOCS2, SOCS3, SOX10, SOX11, SOX2, SOX9, SPEN, SPI1, SPOP, SRC, SRSF2, SSX1, STAG2, STAT3, STAT4, STAT5A, STAT5B, STAT6, STK11, STK36, SUFU, SUZ12, SYK, SYNE1, TAF1, TAF1L, TAL1, TBL1XR1, TBX22, TCF12, TCF3, TCF7L1, TCF7L2, TCL1A, TET1, TET2, TFE3, TGFBR2, TGM7, THBS1, TIMP3, TLL2, TLR4, TLX1, TMEM30A, TMSL3, TNFAIP3, TNFRSF11A, TNFRSF14, TNFRSF17, TNK2, TOP1, TP53, TP63, TPR, TRAF2, TRAF3, TRAF5, TRAF7, TRIM24, TRIM33, TRIP11, TRRAP, TSC1, TSC2, TSHR, TUSC3, TYK2, U2AF1, U2AF2, UBR5, UGT1A1, USP9X, VHL, WAS, WDR90, WHSC1, WISP3, WRN, WT1, XBP1, XPA, XPC, XPO1, XRCC2, YY1AP1, ZBTB16, ZMYM3, ZNF217, ZNF24, ZNF384, ZNF521, ZNF703*, and *ZRSR2*.

**Genes Assessed by Next-Generation Sequencing for Tumor Mutation Burden Determination**

The 432 genes includes: *ABL1, ABL2, ACTB, ACVR1B, AKT1, AKT2, AKT3, ALK, APC, APH1A, AR, ARAF, ARFRP1, ARHGAP26, ARID1A, ARID1B, ARID2, ASMTL, ASXL1, ATM, ATR, ATRX, AURKA, AURKB, AXIN1, AXL, B2M, BAP1, BARD1, BCL10, BCL11B, BCL2, BCL2L1, BCL2L2, BCL6, BCL7A, BCOR, BCORL1, BCR, BIRC3, BLM, BRAF, BRCA1, BRCA2, BRD4, BRIP1, BRSK1, BTG2, BTK, BTLA, CAD, CALR, CARD11, CBFB, CBL, CBLB, CCND1, CCND2, CCND3, CCNE1, CCT6B, CD22, CD274, CD36, CD58, CD70, CD79A, CD79B, CDC73, CDH1, CDK12, CDK4, CDK6, CDK8, CDKN1B, CDKN2A, CDKN2B, CDKN2C, CEBPA, CHD2, CHEK1, CHEK2, CIC, CIITA, CKS1B, CPS1, CREBBP, CRKL, CRLF2, CSF1R, CSF3R, CTCF, CTNNA1, CTNNB1, CUX1, CXCR4, CYLD, DAXX, DDR2, DDX3X, DEK, DICER1, DNM2, DNMT3A, DOT1L, DTX1, DUSP2, DUSP9, EBF1, ECT2L, EED, EGFR, ELP2, EML4, EP300, EPHA3, EPHA5, EPHA7, EPHB1, ERBB2, ERBB3, ERBB4, ERG, ESR1, ETS1, ETV6, EXOSC6, EZH2, FAF1, FAM46C, FANCA, FANCC, FANCD2, FANCE, FANCF, FANCG, FANCL, FAS, FBXO11, FBXO31, FBXW7, FGF10, FGF14, FGF19, FGF23, FGF3, FGF4, FGF6, FGFR1, FGFR2, FGFR3, FGFR4, FH, FHIT, FLCN, FLT1, FLT3, FLT4, FLYWCH1, FOXL2, FOXO1, FOXO3, FOXP1, FRS2, FUBP1, GADD45B, GATA1, GATA2, GATA3, GID4, GNA11, GNA12, GNA13, GNAQ, GNAS, GPR124, GRIN2A, GSK3B, GTSE1, H3F3A, HDAC1, HDAC4, HDAC7, HGF, HIST1H1C, HIST1H1D, HIST1H1E, HIST1H2AC, HIST1H2AG, HIST1H2AL, HIST1H2AM, HIST1H2BC, HIST1H2BJ, HIST1H2BK, HIST1H2BO, HIST1H3B, HNF1A, HRAS, HSP90AA1, ICK, ID3, IDH1, IDH2, IGF1R, IGF2, IKBKE, IKZF1, IKZF2, IKZF3, IL7R, INHBA, INPP4B, INPP5D, IRF1, IRF4, IRF8, IRS2, JAK1, JAK2, JAK3, JARID2, JUN, KAT6A, KDM2B, KDM4C, KDM5A, KDM5C, KDM6A, KDR, KEAP1, KIT, KLF4, KLHL6, KMT2A, KMT2B, KMT2D, KRAS, LEF1, LMO1, LRP1B, LRRK2, MAF, MAFB, MAGED1, MALT1, MAP2K1, MAP2K2, MAP2K4, MAP3K1, MAP3K14, MAP3K6, MAP3K7, MAPK1, MCL1, MDM2, MDM4, MED12, MEF2B, MEF2C, MEN1, MET, MIB1, MITF, MKI67, MLH1, MPL, MRE11A, MSH2, MSH3, MSH6, MTOR, MUTYH, MYC, MYCL1, MYCN, MYD88, MYO18A, NCOR2, NCSTN, NF1, NF2, NFE2L2, NFKBIA, NKX2-1, NLRP1, NOD1, NOTCH1, NOTCH2, NPM1, NRAS, NSD1, NT5C2, NTRK1, NTRK2, NTRK3, NUP93, NUP98, P2RY8, PAG1, PAK3, PALB2, PASK, PAX5, PBRM1, PC, PCBP1, PCLO, PDCD11, PDCD1LG2, PDGFRA, PDGFRB, PDK1, PHF6, PIGA, PIK3C2B, PIK3CA, PIK3CB, PIK3CG, PIK3R1, PIK3R2, PIM1, PLCG2, PML, PMS2, POT1, PPP2R1A, PRDM1, PRKAR1A, PRKDC, PRSS8, PTCH1, PTEN, PTPN11, PTPN2, PTPN6, PTPRO, RAD21, RAD50, RAD51, RAF1, RARA, RASGEF1A, RB1, RECQL4, RELN, RET, RHOA, RHOH, RICTOR, RNF43, ROS1, RPTOR, RUNX1, RUNX1T1, S1PR2, SDHA, SDHB, SDHC, SDHD, SERP2, SETBP1, SETD2, SF3B1, SGK1, SMAD2, SMAD4, SMARCA1, SMARCA4, SMARCB1, SMC1A, SMC3, SMO, SOCS1, SOCS2, SOCS3, SOX10, SOX2, SOX9, SPEN, SPOP, SRC, SRSF2, STAG2, STAT3, STAT4, STAT5A, STAT5B, STAT6, STK11, SUFU, SUZ12, SYK, SYNE1, TAF1, TBL1XR1, TCF3, TCL1A, TET2, TGFBR2, TLL2, TMEM30A, TNFAIP3, TNFRSF11A, TNFRSF14, TNFRSF17, TOP1, TP53, TP63, TRAF2, TRAF3, TRAF5, TRAF7, TSC1, TSC2, TSHR, TUSC3, TYK2, U2AF1, U2AF2, VHL, WDR90, WHSC1, WISP3, WT1, XBP1, XPO1, YY1AP1, ZMYM3, ZNF217, ZNF24, ZNF703*, and *ZRSR2*.

**Tumor Clonality and Cellularity Analysis Formulas**

The cellular representation of a CNV was calculated using the formula ρ=(1-2BAF)/(1-BAF), where ρ represents the percentage of total cells harboring a particular CNV, and BAF is the B-allele frequency. For mutations, variant allele frequencies (z) and percentage of cells harboring a particular CNV (ρ=y) were used to calculate the number of alleles (x) carrying the mutation, which could then be converted to percentage of cells (% cells) harboring the mutation. A few assumptions are necessary to perform this analysis when there are concurrent CNVs. When there is concurrent CN gain, we assume that the additional chromosome also harbors the mutation detected in the original allele so that 2 out of 3 alleles now carry the mutation. When there is concurrent CN loss, we assume that the remaining allele harbors the mutation so that 1 of 1 allele now carry the mutation. When there is CN-LOH, we assume both alleles harbor the mutation so that 2 of 2 alleles now carry the mutation. Example calculations for different scenarios are provided in supplemental **Figures S1-S3**. When there is no concurrent CNV, % cell=x=200z. When there is co-occurring CN gain, x=200z+zy. If x≥2y, % cells=x-y. If x<2y, % cells=x-a where “a” is the largest whole number divisible into x. When there is co-occurring CN loss, x=200z-zy=% cells. If x>=y, % cells = x. If x<y, % cells = x. When there is co-occurring copy-neutral loss-of-heterozygosity (CN-LOH), x=200z. If x≥2y, % cells=x-a where “a” is the largest whole number divisible into x. If x<2y, % cells=x/2. No formula is generated for high-copy gain, or amplification, since there was only one such CNV event, and this did not coincide with DNA mutations.

**Supplemental Results**

**Detailed Pathologic Findings**

In total, 16 cases of LGFL meeting the inclusion and exclusion criteria detailed in Methods were identified from the JHH (N = 5) and NCI (N = 11) pathology archives with the original biopsy procedures performed between 2006 and 2014. All cases underwent pathology consensus review by the study authors. Summary patient and pathologic findings are detailed in **Table 1**. The median/mean age at diagnosis was 53 years with 68.8% of patients being female (N=11) and 31.3% being male (N=5). All patients were alive at the time of data analysis, with a median follow-up of 2.5 years (ranging from 1-9 years). Histologically, all cases showed complete or near-complete nodal architectural effacement, and replacement by a mostly diffuse proliferation of atypical small lymphocytes. Except for one case (case 16), which had approximately 75% diffuse pattern, all other cases showed >75% diffuse growth (**Figure 1** and **Table 1**). Seven cases (7/16; 43.7%) demonstrated focal or partial micro-follicle formation (**Figure 1**). These micro-follicles lacked polarization, mantle zones and tingible-body macrophages, and comprised a mixture of small atypical lymphocytes and occasional larger centroblasts. Background sclerosis and interstitial fibrosis was prominent in 9 cases (9/16; 56.2%). Six cases (6/16; 37.5%) contained residual entrapped normal lymph node with reactive germinal centers. In general, the atypical lymphocytes were small and showed more rounded nuclear cytology (8/16, 50.0%), as opposed to entirely cleaved nuclei characteristic of centrocytes in cFL. Cases with micro-follicles tended to have more centrocyte-like nuclei, which were enriched within micro-follicles, as opposed to more rounded nuclei in cases without micro-follicles. Irrespective of growth pattern, the number of centroblasts per high power field (HPF) was uniformly low (below 15/HPF). Mitotic and apoptotic figures were only rarely identified.

Routine immunohistochemical staining showed that all the lymphomas expressed CD10, BCL6 and CD23 (**Figure 1** and **Table 1**), although staining patterns varied in intensity (weak, variable, strong) and proportion of stained cells (focal, patchy and diffuse). Although 5 of 16 cases (from JHH) were selected using CD23 as an inclusion criteria, the remaining 11 cases (from NCI) were not selected based on CD23 status, yet all cases demonstrated consistent CD23 expression. Some degree of BCL2 positivity was also identified in 13 of 16 cases (81.2%). Two cases showed equivocal BCL2 staining (Cases 7 and 8) due to the extensive T-cell infiltrate, and one case was BCL2 negative (Case 15). Two cases with a micro-follicular growth pattern showed disparate BCL2 expression (case 6 and 9) where the diffuse area demonstrated weak-positive and positive expression, respectively, and the micro-follicles were either equivocal or lacked BCL2 expression, respectively. Of note, case 9 also showed disparate staining patterns in the diffuse versus micro-follicle components for CD10 (patchy and weak positive, respectively) and CD23 (positive and negative, respectively). Follicular dendritic cell networks, as demonstrated by either CD21 and/or CD23, were absent in all cases, or only focally intact, and were not co-localized with micro-follicles. Finally, the overall Ki-67 proliferation index, when available (10/16 cases), was low, averaging approximately 15-20%. Three cases with micro-follicles showed marginally higher Ki-67 proliferation rates in the micro-follicles (Cases 3, 9, 14). Based on immunohistochemical stains, neoplastic B-cells comprised at least 20% of cells in all cases, and the background T-cell infiltrates ranged from moderate to extensive.

**Detailed Next-Generation Sequencing Findings**

A median of 43,798,939 unique reads (ranging from 25,439,236- 61,825,336) were mapped to the human genome per case. The median PCR duplication rate was 43.9% (ranging from 23.8–81.3%), and the 2 low quality samples showed PCR duplication rates of 59.6% and 81.3%, respectively. The median unique depth of coverage was 570x (ranging from 157–853x) with a median of 99.2% (ranging from 96.6-99.5%) panel specific coverage of ≥150x. The two low quality samples showed unique depth of coverage values of 157x and 171x, respectively.

**Detailed Clonality/Tumor Purity Analysis Findings**

Integrating both mutational VAFs and (co-occurring) CNV B-allele frequencies, the cellular representation of individual alterations was calculated, which enabled estimation of tumor purity/content. The calculated tumor purity ranged from 15.1-87.2% with a median of 57.1% (95% CI of 33.3-79.8%). The case with no detectable CNVs (case 12) had the lowest estimated tumor cellularity (15.1%) based on clonality analysis. Tumor purity of lymphomas with (mean 52.1%, median 46.1%) and without (mean 74.4%, median 79.8%) *CREBBP* and *STAT6* co-mutations were not statistically significantly different. CNVs involving 16p13.3 accounted for 27.2-100% of tumor cells (median of 72.5%, 95 CI 39.4-86.7%), while CNVs involving 1p36.3 accounted for 49.5-86.7% of tumor cells (median of 69.2%, 95 CI 49.5-86.7%).

**Detailed Statistical Analysis Findings**

The threshold for statistical significance was set at a p-value of ≤ 0.05. When appropriate, median values and 95% confidence intervals around the median were calculated. When continuous variables of total detected alterations (**Figure 4**) and tumor cellularity/purity (**Figure 6**) were compared using a t-test, several tests for normality were also performed, including Anderson-Darling (A2*), D'Agostino-Pearson omnibus (K2), Shapiro-Wilk (W) and Kolmogorov-Smirnov (distance), which all confirmed that these subgroup comparisons derived from normal distributions with an alpha error of 0.05. The QQ plots are provided in supplemental **Figure S4**. The two groups showed different standard error of mean with respect to total detected alterations (0.8385 for cases with *CREBBP/STAT6* co-mutations and 3.930 for those without), but showed similar standard error of mean values with respective to tumor cellularity/purity (6.309 for cases with *CREBBP/STAT6* co-mutations and 8.821 for those without).

**Data and Method Sharing**

All SNP-microarray data and next-generation sequencing data will be submitted to a public data repository, and the accession ID will be provided when assigned. All bioinformatics tools used for this study are publically available, except for MDLVC, which is a custom variant caller pipeline designed for clinical use in the Johns Hopkins Molecular Diagnostic Laboratory. As this is a clinical tool, we are unable to publically share the source code. However, interested investigators are encouraged to contact the corresponding author for access to the source code.

**Supplemental Figure and Table Legends**

**Figure S1. Example tumor cellularity estimation when there are no concurrent CNVs, and when there is a concurrent CN gain.** Tumor cells are displayed as blue circles, while normal cells are represented as cartoon neutrophils. The red “X” indicates a mutation detected by sequencing.

**Figure S2. Example tumor cellularity estimation when there is concurrent CN loss.** Tumor cells are displayed as blue circles, while normal cells are represented as cartoon neutrophils. The red “X” indicates a mutation detected by sequencing.

**Figure S3. Example tumor cellularity estimation when there is concurrent CN-LOH.** Tumor cells are displayed as blue circles, while normal cells are represented as cartoon neutrophils. The red “X” indicates a mutation detected by sequencing. Yellow shading denotes CN-LOH.

**Figure S4. QQ plots for tests of normal distribution in subgroup analyses.** **A**. QQ plot for total detected alterations in cases with *CREBBP/STAT6* co-mutations and those without. **B**. QQ plot for tumor cellularity in cases with *CREBBP/STAT6* co-mutations and those without.

**Table S1. Complete List of Detected CNVs.** All CNVs are annotated with the affected chromosome region, cytoband, CNV event (loss, gain, CN-LOH), length of the change in base pairs, and the number of genes contained in each region.

**Table S2. Complete List of Detected Mutations**. All mutations are annotated with the chromosomal coordinate, base change, affected region, gene name, total alternative and reference reads, calculated VAF, cDNA and protein change with transcript, and functional consequence.

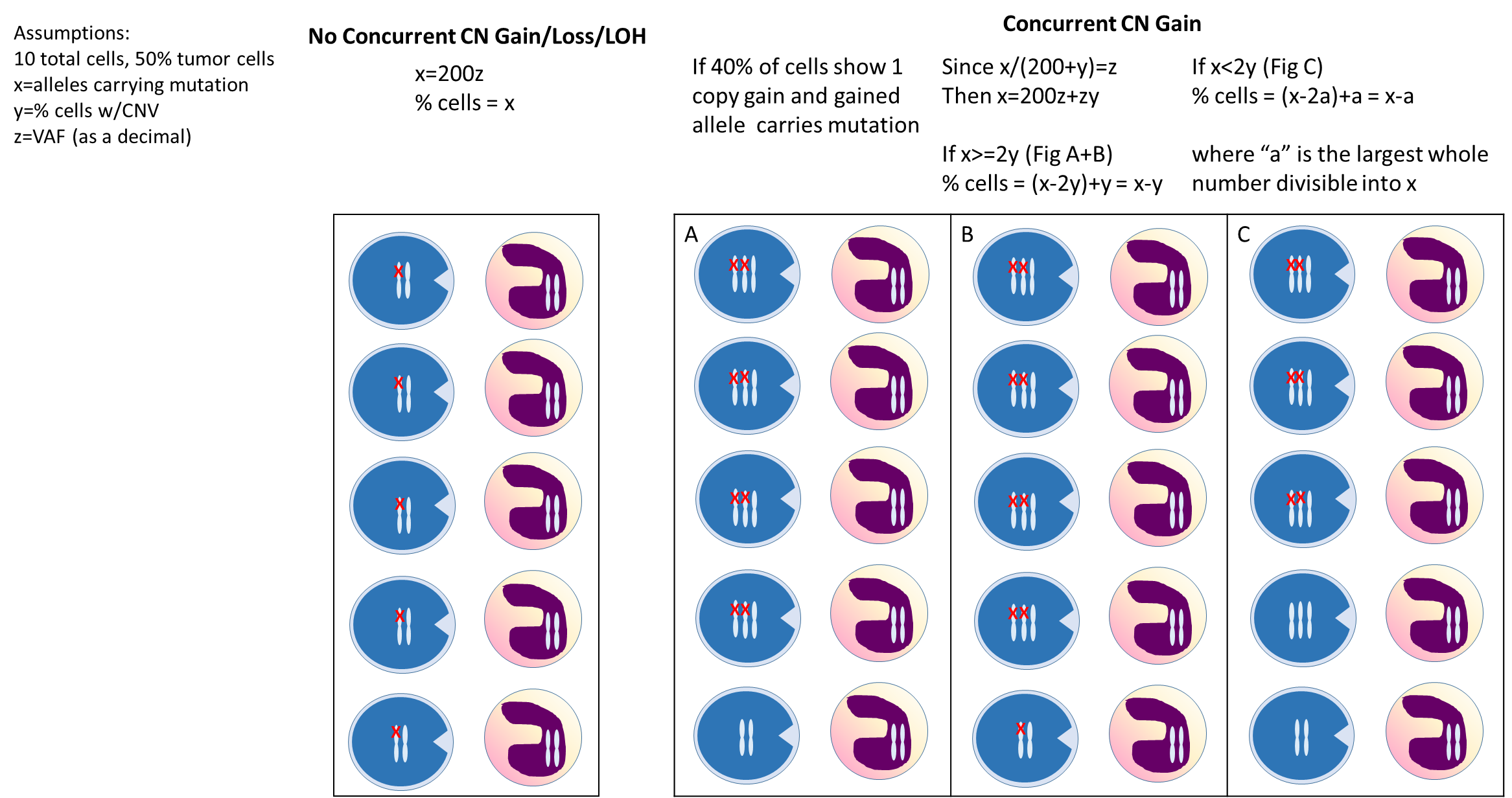
**Figure S1**

**Figure S2**

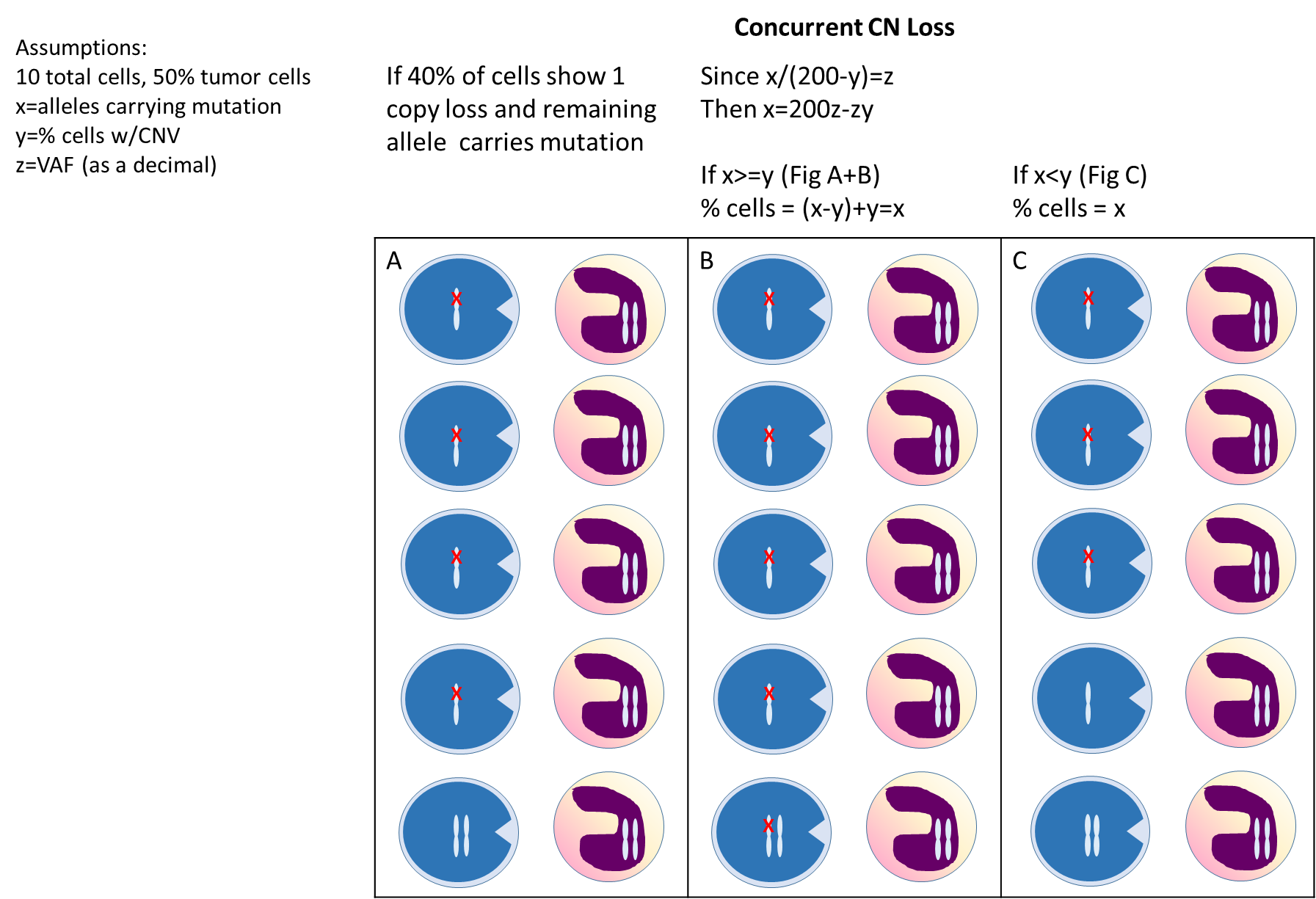

**Figure S3**

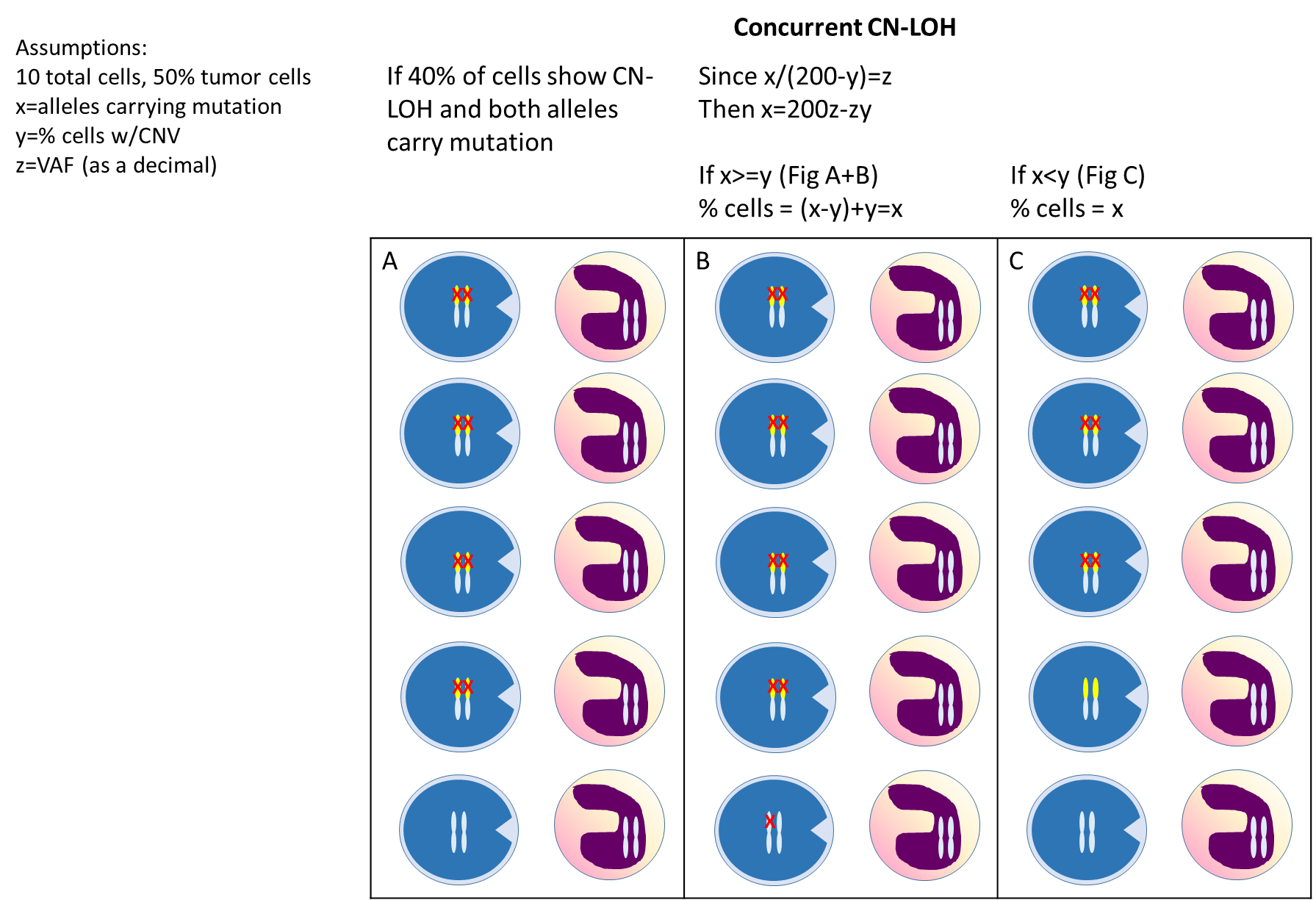

**Figure S4**

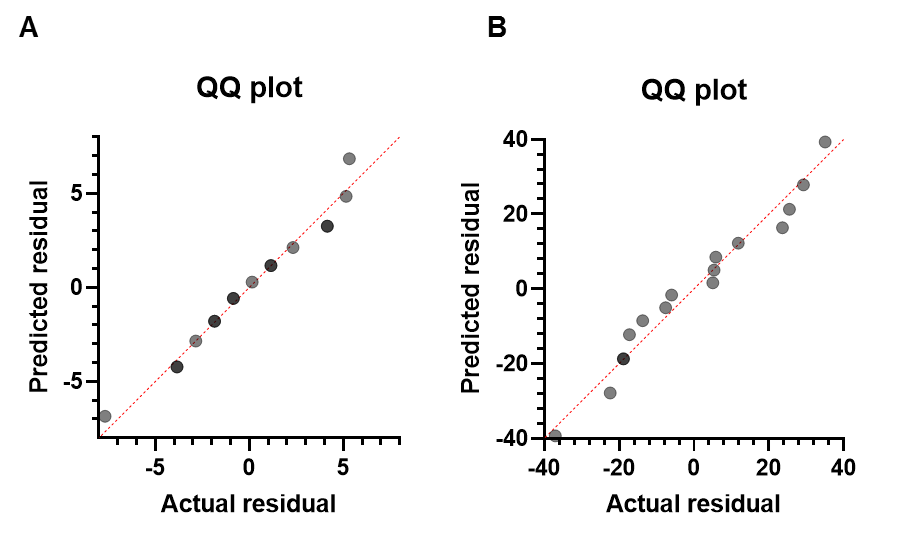

**Table S1. Complete List of Detected CNVs.**

| Case | Chromosome Region | Cytoband | CNV Event | CNV Length (bp) | CNV Length (Mb) | No. of Genes in Region |
| --- | --- | --- | --- | --- | --- | --- |
| 1 | chr1:1-111,449,999 | p36.33 - p13.3 | CN-LOH | 111449999 | 111.45 | 1282 |
| 1 | chr16:1-30,408,765 | p13.3 - p11.2 | Loss | 30156748 | 30.16 | 519 |
| 2 | chr1:1-38,243,785 | p36.33 - p34.3 | CN-LOH | 111449999 | 111.45 | 1282 |
| 2 | chr2:1-242,421,866 | p25.3 - q37.3 | Gain | 30156748 | 30.16 | 519 |
| 2 | chr12:49,149,597-133,764,292 | q13.12 - q24.33 | Gain | 84614696 | 84.61 | 861 |
| 2 | chr16:1-30,156,748 | p13.3 - p11.2 | Loss | 30156748 | 30.16 | 519 |
| 3 | chr2:47,896,026-65,713,066 | p16.3 - p14 | High Copy Gain | 17817041 | 17.82 | 86 |
| 3 | chr5:1-180,915,260 | p15.33 - q35.3 | Gain | 2941529 | 2.94 | 14 |
| 3 | chr6:0-57,526,313 | p25.3 - p11.2 | Gain | 2941529 | 2.94 | 14 |
| 3 | chr6:61,000,000-171,115,067 | q11.1 - q27 | Loss | 2941529 | 2.94 | 14 |
| 3 | chr8:122,844,682-146,364,022 | q24.13 - q24.3 | Gain | 2941529 | 2.94 | 14 |
| 3 | chr9:74,477,270-75,685,931 | q21.13 | Loss | 2941529 | 2.94 | 14 |
| 3 | chr12:1-122,301,299 | p13.33 - q24.31 | Gain | 111449999 | 111.45 | 1282 |
| 3 | chr12:122,346,052-133,851,895 | q24.31 - q24.33 | CN-LOH | 111449999 | 111.45 | 1282 |
| 3 | chr15:43,486,861-102,148,164 | q15.2 - q26.3 | CN-LOH | 84614696 | 84.61 | 861 |
| 4 | chr1:1-31,308,177 | p36.33 - p35.2 | CN-LOH | 111449999 | 111.45 | 1282 |
| 4 | chr16:1-31,151,521 | p13.3 - p11.2 | CN-LOH | 30156748 | 30.16 | 519 |
| 5 | chr3:109,663-10,199,184 | p26.3 - p25.3 | Loss | 17817041 | 17.82 | 86 |
| 5 | chr15:42,438,564-102,531,392 | q15.1 - q26.3 | CN-LOH | 84614696 | 84.61 | 861 |
| 6 | chr4:178,147,903-190,760,143 | q34.3 - q35.2 | Loss | 2941529 | 2.94 | 14 |
| 6 | chr6:1-42,686,408 | p25.3 - p21.1 | CN-LOH | 2941529 | 2.94 | 14 |
| 6 | chr8:1-146,168,128 | p23.3 - q24.3 | Gain | 2941529 | 2.94 | 14 |
| 6 | chr16:1-14,146,098 | p13.3 - p13.12 | Loss | 84614696 | 84.61 | 861 |
| 6 | chrX:115,453,556-155,233,098 | q23 - q28 | Gain | 2941529 | 2.94 | 14 |
| 7 | chr16:1-18,143,302 | p13.3 - p12.3 | Loss | 84614696 | 84.61 | 861 |
| 8 | chr6:81,382,561-96,729,995 | q14.1 - q16.1 | Loss | 2941529 | 2.94 | 14 |
| 8 | chr8:1-145,802,447 | p23.3 - q24.3 | Gain | 2941529 | 2.94 | 14 |
| 9 | chr1:828,472-5,928,631 | p36.33 - p36.31 | Loss | 111449999 | 111.45 | 1282 |
| 9 | chr6:1-30,989,385 | p25.3 - p21.33 | Loss | 2941529 | 2.94 | 14 |
| 9 | chr8:1-146,288,066 | p23.3 - q24.3 | Gain | 2941529 | 2.94 | 14 |
| 10 | chr16:1-14,750,607 | p13.3 - p13.12 | Loss | 84614696 | 84.61 | 861 |
| 11 | chr1:1-31,878,470 | p36.33 - p35.2 | Loss | 111449999 | 111.45 | 1282 |
| 11 | chr1:145,563,516-248,578,622 | q21.1 - q44 | Gain | 111449999 | 111.45 | 1282 |
| 11 | chr8:13,502,822-14,759,867 | p22 | Gain | 2941529 | 2.94 | 14 |
| 11 | chr8:48,916,052-146,128,340 | q11.21 - q24.3 | CN-LOH | 2941529 | 2.94 | 14 |
| 11 | chr14:84,668,767-106,004,323 | q31.2 - q32.33 | CN-LOH | 84614696 | 84.61 | 861 |
| 11 | chr16:1-31,855,686 | p13.3 - p11.2 | Loss | 30156748 | 30.16 | 519 |
| 13 | chr1:876,437-15,425,146 | p36.33 - p36.21 | Loss | 111449999 | 111.45 | 1282 |
| 13 | chr2:203,572,362-243,199,373 | q33.2 - q37.3 | Gain | 30156748 | 30.16 | 519 |
| 13 | chr16:1-7,146,126 | p13.3 | Loss | 30156748 | 30.16 | 519 |
| 13 | chr18:70,042,133-74,879,435 | q22.3 - q23 | Loss | 30156748 | 30.16 | 519 |
| 13 | chr18:75,135,720-78,077,248 | q23 | Gain | 30156748 | 30.16 | 519 |
| 14 | chr11:1-51,197,691 | p15.5 - p11.12 | Gain | 111449999 | 111.45 | 1282 |
| 14 | chr16:222,778-31,656,289 | p13.3 - p11.2 | Loss | 30156748 | 30.16 | 519 |
| 15 | chr1:1-32,404,477 | p36.33 - p35.1 | Loss | 111449999 | 111.45 | 1282 |
| 15 | chr2:95,537,000-243,199,373 | q11.1 - q37.3 | Gain | 17817041 | 17.82 | 86 |
| 15 | chr16:0-16,685,044 | p13.3 - p13.11 | Loss | 84614696 | 84.61 | 861 |
| 15 | chr16:16,685,044-35,069,526 | p13.11 - p11.1 | Gain | 30156748 | 30.16 | 519 |
| 15 | chr16:48,589,126-90,354,753 | q12.1 - q24.3 | Gain | 30156748 | 30.16 | 519 |
| 16 | NA | NA | NA | NA | NA | NA |

**Table S2. Complete List of Detected Mutations**.

| Case | Chr:Pos | Base Change | Gene | Alt+,Alt-/Ref+,Ref- | %VAF | cDNA change | Protein change | Consequence |
| --- | --- | --- | --- | --- | --- | --- | --- | --- |
| 1 | chr16:3781323 | AAGG>A | *CREBBP* | 33,49/131,143 | 23.03 | NM_004380.2:c.5039_5041del | NP_004371.2:p.Ser1680del | inframe deletion |
| 1 | chr6:26157013 | C>T | *HIST1H1E* | 17,26/191,183 | 10.31 | NM_005321.2:c.395C>T | NP_005312.1:p.Ala132Val | missense |
| 1 | chr12:49420274 | G>A | *KMT2D* | 79,44/556,396 | 11.44 | NM_003482.3:c.15475C>T | NP_003473.3:p.Arg5159Trp | missense |
| 1 | chr19:19260226 | T>G | *MEF2B* | 5,11/65,115 | 8.16 | NM_001145785.1:c.67A>C | NP_001139257.1:p.Lys23Gln | missense |
| 1 | chr1:144854167 | CTAAGTTAGTCCTT>C | *PDE4DIP* | 89,73/1349,1047 | 6.33 | NM_014644.5:c.6984_6996del | NP_055459.5:p.Thr2330Ter | stop_gained |
| 1 | chr12:57496662 | C>G | *STAT6* | 35,37/194,213 | 15.03 | NM_003153.4:c.1255G>C | NP_003144.3:p.Asp419His | missense |
| 1 | chr1:2489220 | G>A | *TNFRSF14* | 31,33/125,179 | 17.39 | NM_003820.2:c.125G>A | NP_003811.2:p.Cys42Tyr | missense |
| 2 | chr16:3789628 | C>T | *CREBBP* | 191,137/178,121 | 52.31 | NM_004380.2:c.4231G>A | NP_004371.2:p.Gly1411Arg | missense |
| 2 | chr7:148508727 | T>G | *EZH2* | 114,49/323,166 | 25 | NM_004456.4:c.1937A>C | NP_004447.2:p.Tyr646Ser | missense |
| 2 | chr13:41240288 | C>T | *FOXO1* | 6,18/14,21 | 40.68 | NM_002015.3:c.62G>A | NP_002006.2:p.Arg21His | missense |
| 2 | chr12:48189070 | C>T | *HDAC7* | 13,7/95,93 | 9.62 | NM_015401.3:c.1181G>A | NP_056216.2:p.Arg394Gln | missense |
| 2 | chr12:49420607 | G>A | *KMT2D* | 68,64/226,244 | 21.93 | NM_003482.3:c.15142C>T | NP_003473.3:p.Arg5048Cys | missense |
| 2 | chr12:49427954 | G>A | *KMT2D* | 65,104/281,309 | 22.27 | NM_003482.3:c.10636C>T | NP_003473.3:p.Gln3546Ter | stop gain |
| 2 | chr16:11348945 | G>A | *SOCS1* | 33,33/98,93 | 25.68 | NM_003745.1:c.391C>T | NP_003736.1:p.Gln131Ter | stop gain |
| 2 | chr16:11349289 | G>A | *SOCS1* | 22,20/66,50 | 26.58 | NM_003745.1:c.47C>T | NP_003736.1:p.Ala16Val | missense |
| 2 | chr12:57496662 | C>T | *STAT6* | 33,59/217,294 | 15.26 | NM_003153.4:c.1255G>A | NP_003144.3:p.Asp419Asn | missense |
| 2 | chr14:96180283 | C>T | *TCL1A* | 59,33/157,86 | 27.46 | NM_021966.2:c.120+1G>A | p.? | splice_donor_variant |
| 2 | chr1:2492067 | CGA>C | *TNFRSF14* | 80,39/90,42 | 47.41 | NM_003820.2:c.469_470del | NP_003811.2:p.Gln158GlyfsTer75 | frameshift |
| 3 | chr15:45003745 | A>G | *B2M* | 25,16/163,92 | 13.85 | NM_001243186.1:c.384G>C | NP_001230115.1:p.Gln128His | missense |
| 3 | chr15:45007645 | C>CA | *B2M* | 128,90/216,150 | 37.33 | NM_001243186.1:c.676G>A | NP_001230115.1:p.Glu226Lys | missense |
| 3 | chr6:70071082 | G>C | *BAI3* | 52,57/26,23 | 68.99 | NM_004048.2:c.93dup | NP_004039.1:p.Arg32ThrfsTer25 | frameshift |
| 3 | chr12:122460051 | C>G | *BCL7A* | 8,14/17,45 | 26.19 | NM_024007.3:c.625C>T | NP_076870.1:p.Arg209Trp | missense |
| 3 | chr17:62007560 | C>T | *CD79B* | 91,86/101,81 | 49.3 | NM_198124.1:c.4416C>A | NP_937757.1:p.Ser1472Arg | missense |
| 3 | chr16:3789597 | C>T | *CREBBP* | 51,33/38,26 | 56.76 | NM_001704.2:c.3917G>C | NP_001695.1:p.Ser1306Thr | missense |
| 3 | chr8:113562928 | G>T | *CSMD3* | 26,15/90,45 | 23.3 | NM_004380.2:c.4262G>A | NP_004371.2:p.Cys1421Tyr | missense |
| 3 | chr8:113694686 | T>C | *CSMD3* | 49,31/125,71 | 28.99 | NM_000626.2:c.304G>A | NP_000617.1:p.Glu102Lys | missense |
| 3 | chr5:138261079 | G>A | *CTNNA1* | 29,43/178,185 | 16.55 | NM_000043.4:c.684_688del | NP_000034.1:p.Asp228GlufsTer2 | frameshift |
| 3 | chr5:158267048 | G>A | *EBF1* | 40,30/250,204 | 13.36 | NM_012465.3:c.1538G>A | NP_036597.1:p.Arg513Lys | missense |
| 3 | chr5:158524085 | C>T | *EBF1* | 76,101/122,125 | 41.75 | NM_020993.3:c.54C>G | NP_066273.1:p.Ile18Met | missense |
| 3 | chr22:41527493 | A>G | *EP300* | 88,88/88,90 | 49.72 | NM_198124.1:c.2542A>G | NP_937757.1:p.Ile848Val | missense |
| 3 | chr10:90773878 | GTTGAC>G | *FAS* | 30,21/107,57 | 23.72 | NM_001429.3:c.1384A>G | NP_001420.2:p.Ser462Gly | missense |
| 3 | chr12:49433060 | G>A | *KMT2D* | 53,70/79,126 | 37.5 | NM_024007.3:c.188G>A | NP_076870.1:p.Arg63Gln | missense |
| 3 | chr1:11307911 | A>T | *MTOR* | 59,66/93,84 | 41.39 | NM_004958.3:c.1081T>A | NP_004949.1:p.Cys361Ser | missense |
| 3 | chr6:37138577 | G>C | *PIM1* | 32,21/76,63 | 27.6 | NM_003482.3:c.8311C>T | NP_003473.3:p.Arg2771Ter | stop gain |
| 3 | chr6:37139063 | G>A | *PIM1* | 26,28/112,122 | 18.75 | NM_004048.2:c.1A>G | NP_004039.1:p.Met1? | start lost |
| 3 | chr10:98155132 | C>T | *TLL2* | 22,40/54,108 | 27.68 | NM_001903.2:c.1882G>A | NM_001903.2(CTNNA1_i001):p.(Ala628Thr) | missense |
| 4 | chr13:32907410 | T>A | *BRCA2* | 20,14/199,140 | 9.12 | NM_000059.3:c.1795T>A | NP_000050.2:p.Ser599Thr | missense |
| 4 | chr16:3788606 | A>T | *CREBBP* | 52,34/182,174 | 19.46 | NM_004380.2:c.4348T>A | NP_004371.2:p.Tyr1450Asn | missense |
| 4 | chr13:41240288 | C>T | *FOXO1* | 8,14/54,84 | 13.75 | NM_002015.3:c.62G>A | NP_002006.2:p.Arg21His | missense |
| 4 | chr6:26157108 | G>C | *HIST1H1E* | 13,27/185,190 | 9.64 | NM_005321.2:c.490G>C | NP_005312.1:p.Ala164Pro | missense |
| 4 | chr12:49432003 | C>A | *KMT2D* | 17,12/177,170 | 7.71 | NM_003482.3:c.9136G>T | NP_003473.3:p.Glu3046Ter | stop gain |
| 4 | chr12:57496661 | T>C | *STAT6* | 7,16/241,222 | 4.73 | NM_003153.4:c.1256A>G | NP_003144.3:p.Asp419Gly | missense |
| 4 | chr1:2488105 | T>A | *TNFRSF14* | 34,23/126,88 | 21.03 | NM_003820.2:c.2T>A | NP_003811.2:p.Met1? | start lost |
| 5 | chr15:45003785 | CTCTT>C | *B2M* | 43,24/94,52 | 31.46 | NM_003820.2:c.218C>T | NP_003811.2:p.Thr73Ile | missense |
| 5 | chr16:3790409 | A>C | *CREBBP* | 20,14/57,41 | 25.76 | NM_003820.2:c.412T>G | NP_003811.2:p.Cys138Gly | missense |
| 5 | chr12:57496661 | T>C | *STAT6* | 43,19/265,134 | 13.45 | NM_003153.4:c.1256A>G | NP_003144.3:p.Asp419Gly | missense |
| 5 | chr1:2489821 | C>T | *TNFRSF14* | 5,9/40,67 | 11.57 | NM_004048.2:c.45_48del | NP_004039.1:p.Ser16AlafsTer27 | frameshift |
| 5 | chr1:2491369 | T>G | *TNFRSF14* | 10,13/53,55 | 17.56 | NM_004380.2:c.4124T>G | NP_004371.2:p.Met1375Arg | missense |
| 5 | chr3:189587161 | G>T | *TP63* | 11,11/31,36 | 24.72 | NM_003722.4:c.1178G>T | NP_003713.3:p.Arg393Leu | missense |
| 6 | chr22:23523243 | G>C | *BCR* | 23,15/152,118 | 12.34 | NM_002015.3:c.459C>G | NP_002006.2:p.Ser153Arg | missense |
| 6 | chr22:23523415 | C>T | *BCR* | 7,17/50,99 | 13.87 | NM_002015.3:c.158A>C | NP_002006.2:p.Asp53Ala | missense |
| 6 | chr22:23523461 | CC>AG | *BCR* | 4,8/27,64 | 11.65 | NM_002015.3:c.70A>G | NP_002006.2:p.Thr24Ala | missense |
| 6 | chr16:3817719 | A>C | *CREBBP* | 19,16/185,88 | 11.36 | NM_004380 :c.3250+2T>G | p.? | splice_donor_variant |
| 6 | chr16:3817768 | T>C | *CREBBP* | 26,22/199,150 | 12.09 | NM_004380.2:c.3196A>G | NP_004371.2:p.Ser1066Gly | missense |
| 6 | chr16:3817775 | T>C | *CREBBP* | 28,23/201,152 | 12.62 | NM_004456.4:c.1936T>A | NP_004447.2:p.Tyr646Asn | missense |
| 6 | chr7:148508728 | A>T | *EZH2* | 26,20/235,147 | 10.75 | NM_003153.4:c.1255G>C | NP_003144.3:p.Asp419His | missense |
| 6 | chr13:41239891 | G>C | *FOXO1* | 4,18/32,90 | 15.28 | NM_004327.3:c.268C>T | NP_004318.3:p.Pro90Ser | missense |
| 6 | chr13:41240192 | T>G | *FOXO1* | 12,14/59,42 | 20.47 | NM_004327.3:c.314_315delinsAG | NM_004327.3(BCR_i001):p.(Ala105Glu) | missense |
| 6 | chr13:41240280 | T>C | *FOXO1* | 8,8/38,53 | 14.95 | NM_004380.2:c.3250+2T>G | p.? | splice_donor_variant |
| 6 | chr12:57496662 | C>G | *STAT6* | 32,32/217,176 | 14 | NM_004327.3:c.96G>C | NP_004318.3:p.Glu32Asp | missense |
| 7 | chr7:2976713 | G>C | *CARD11* | 13,4/200,94 | 5.47 | NM_032415.4:c.1299C>G | NP_115791.3:p.Ser433Arg | missense |
| 7 | chr2:208420447 | C>T | *CREB1* | 50,47/317,314 | 13.32 | NM_134442.3:c.88C>T | NP_604391.1:p.Gln30Ter | stop gain |
| 7 | chr16:3788606 | A>T | *CREBBP* | 36,32/198,184 | 15.11 | NM_004380.2:c.4348T>A | NP_004371.2:p.Tyr1450Asn | missense |
| 7 | chr22:41536160 | C>T | *EP300* | 20,15/348,244 | 5.58 | NM_001429.3:c.1777C>T | NP_001420.2:p.Pro593Ser | missense |
| 7 | chr16:15831423 | T>G | *MYH11* | 16,18/107,121 | 12.98 | NM_022844.2:c.3176A>C | NP_074035.1:p.Lys1059Thr | missense |
| 7 | chr7:82580213 | A>G | *PCLO* | 41,37/389,424 | 8.75 | NM_033026.5:c.9691T>C | NP_149015.2:p.Phe3231Leu | missense |
| 7 | chr12:57496662 | C>T | *STAT6* | 16,17/185,195 | 7.99 | NM_003153.4:c.1255G>C | NP_003144.3:p.Asp419His | missense |
| 7 | chr1:2489265 | G>A | *TNFRSF14* | 13,23/72,134 | 14.88 | NM_003820.2:c.170G>A | NP_003811.2:p.Cys57Tyr | missense |
| 7 | chr17:7577538 | C>T | *TP53* | 14,7/137,101 | 8.11 | NM_000546.5:c.743G>A | NM_000546.5(TP53_i001):p.(Arg248Gln) | missense |
| 8 | chr1:2489234 | T>A | *TNFRSF14* | 8,17/104,155 | 8.8 | NM_025003.3:c.3229T>C | NP_079279.3:p.Cys1077Arg | missense |
| 8 | chr1:65351944 | G>A | *JAK1* | 152,169/196,204 | 44.52 | NM_004380.2:c.5039_5041del | NP_004371.2:p.Ser1680del | inframe deletion |
| 8 | chr3:47098561 | G>A | *SETD2* | 71,81/90,99 | 44.57 | NM_003745.1:c.16C>T | NP_003736.1:p.Gln6Ter | stop_gained |
| 8 | chr7:148508728 | A>T | *EZH2* | 57,27/352,197 | 13.27 | NM_003153.4:c.1114G>A | NP_003144.3:p.Glu372Lys | missense |
| 8 | chr12:43825167 | A>G | *ADAMTS20* | 75,51/606,442 | 10.73 | NM_004456.4:c.1936T>A | NP_004447.2:p.Tyr646Asn | missense |
| 8 | chr12:57498345 | C>T | *STAT6* | 14,25/95,86 | 17.73 | NM_002227.2:c.4C>T | NP_002218.2:p.Gln2Ter | stop_gained |
| 8 | chr16:11348794 | C>G | *SOCS1* | 28,40/159,201 | 15.89 | NM_021140.2:c.1177C>T | NP_066963.2:p.Arg393Ter | stop_gained |
| 8 | chr16:11349320 | G>A | *SOCS1* | 29,28/133,157 | 16.43 | NM_014159.6:c.6713C>T | NP_054878.5:p.Ser2238Phe | missense |
| 8 | chr16:3781323 | AAGG>A | *CREBBP* | 31,25/142,159 | 15.69 | NM_004380.2:c.5039_5041del | NP_004371.2:p.Ser1680del | inframe deletion |
| 8 | chrX:44918694 | C>T | *KDM6A* | 7,34/136,410 | 6.98 | NM_003820.3:c.139T>A | NP_003811.2:p.Tyr47Asn | missense |
| 9 | chr1:2491394 | CG>C | *TNFRSF14* | 27,55/74,149 | 26.89 | NM_032415.4:c.3304G>A | NP_115791.3:p.Val1102Met | missense |
| 9 | chr2:202149998 | A>G | *CASP8* | 56,71/188,312 | 20.26 | NM_004456.4:c.1936T>A | NP_004447.2:p.Tyr646Asn | missense |
| 9 | chr2:61720140 | C>T | *XPO1* | 106,109/165,159 | 39.89 | NM_005321.2:c.307_308delinsAA | NP_005312.1:p.Gly103Asn | missense |
| 9 | chr6:26156926 | G>A | *HIST1H1E* | 41,76/95,140 | 33.24 | NM_003153.4:c.1256A>G | NP_003144.3:p.Asp419Gly | missense |
| 9 | chr7:148508728 | A>T | *EZH2* | 77,45/267,162 | 22.14 | NM_003820.2:c.440del | NP_003811.2:p.Gly147AlafsTer43 | frameshift |
| 9 | chr7:2946433 | C>T | *CARD11* | 48,61/169,217 | 22.02 | NM_003400.3:c.1294G>A | NP_003391.1:p.Val432Ile | missense |
| 9 | chr12:57496661 | T>C | *STAT6* | 91,84/302,322 | 21.9 | NM_033356.3:c.1217A>G | NP_203520.1:p.Tyr406Cys | missense |
| 9 | chr17:78341820 | C>A | *RNF213* | 67,58/294,282 | 17.83 | NM_031891.2:c.1873A>T | NP_114097.2:p.Ile625Phe | missense |
| 9 | chr18:59217435 | A>T | *CDH20* | 23,32/113,149 | 17.35 | NM_001429.3:c.4198A>T | NP_001420.2:p.Ser1400Cys | missense |
| 9 | chr22:23523232 | G>A | *BCR* | 54,36/180,146 | 21.63 | NM_001256071.1:c.12032C>A | NP_001243000.1:p.Pro4011His | missense |
| 9 | chr22:41565532 | A>T | *EP300* | 36,42/185,148 | 18.98 | NM_015001.2:c.2692A>G | NP_055816.2:p.Lys898Glu | missense |
| 10 | chr1:65307173 | G>A | *JAK1* | 49,42/445,425 | 9.47 | NM_002227.3:c.2515C>T | NP_002218.2:p.Arg839Ter | stop gain |
| 10 | chr7:2979513 | A>G | *CARD11* | 38,44/455,501 | 7.9 | NM_032415.4:c.734T>C | NP_115791.3:p.Leu245Pro | missense |
| 10 | chr7:82544458 | G>T | *PCLO* | 66,55/582,545 | 9.7 | NM_033026.5:c.12844C>A | NP_149015.2:p.Pro4282Thr | missense |
| 10 | chr12:57496671 | C>G | *STAT6* | 46,57/368,412 | 11.66 | NM_003153.4:c.1246G>C | NP_003144.3:p.Gly416Arg | missense |
| 10 | chr13:41240349 | T>A | *FOXO1* | 2,5/24,38 | 10.14 | NM_002015.3:c.1A>T | NP_002006.2:p.Met1? | start lost |
| 10 | chr16:11348998 | A>G | *SOCS1* | 21,21/182,217 | 9.52 | NM_003745.1:c.338T>C | NP_003736.1:p.Phe113Ser | missense |
| 10 | chr16:11349109 | G>A | *SOCS1* | 8,16/64,99 | 12.83 | NM_003745.1:c.227C>T | NP_003736.1:p.Ala76Val | missense |
| 10 | chr16:11349134 | T>C | *SOCS1* | 8,10/71,85 | 10.34 | NM_003745.1:c.202A>G | NP_003736.1:p.Thr68Ala | missense |
| 10 | chr16:11349146 | A>T | *SOCS1* | 9,10/66,79 | 11.59 | NM_003745.1:c.190T>A | NP_003736.1:p.Tyr64Asn | missense |
| 10 | chr16:3789596 | G>C | *CREBBP* | 79,52/362,230 | 18.12 | NM_004380.2:c.4263C>G | NP_004371.2:p.Cys1421Trp | missense |
| 11 | chr1:2491376 | CT>C | *TNFRSF14* | 67,98/173,262 | 27.5 | NM_003820.2:c.421del | NP_003811.2:p.Tyr141ThrfsTer49 | frame shift |
| 11 | chr5:38512036 | C>T | *LIFR* | 28,64/255,412 | 12.12 | NM_002310.5:c.592G>A | NP_002301.1:p.Asp198Asn | missense |
| 11 | chr7:5568811 | T>C | *ACTB* | 44,36/317,186 | 13.72 | NM_001101.3:c.344A>G | NP_001092.1:p.Asn115Ser | missense |
| 11 | chr7:5568824 | T>G | *ACTB* | 45,34/347,192 | 12.78 | NM_001101.3:c.331A>C | NP_001092.1:p.Asn111His | missense |
| 11 | chr7:5568938 | G>A | *ACTB* | 38,41/377,427 | 8.95 | NM_001101.3:c.217C>T | NP_001092.1:p.His73Tyr | missense |
| 11 | chr7:80276129 | A>C | *CD36* | 98,103/575,705 | 13.57 | ENSESTT00000004568.1:c.73A>C | ENSESTP00000004568.1:p.Ile25Leu | missense |
| 11 | chr7:82538261 | G>A | *PCLO* | 89,78/703,580 | 11.52 | NM_033026.5:c.13369C>T | NP_149015.2:p.His4457Tyr | missense |
| 11 | chr7:82583358 | G>T | *PCLO* | 102,115/707,743 | 13.02 | NM_033026.5:c.6911C>A | NP_149015.2:p.Ala2304Asp | missense |
| 11 | chr8:93017520 | G>T | *RUNX1T1* | 36,92/167,348 | 19.91 | NM_001198627.1:c.564C>A | NP_001185556.1:p.Asn188Lys | missense |
| 11 | chr13:41240288 | C>G | *FOXO1* | 8,10/46,80 | 12.5 | NM_002015.3:c.62G>C | NP_002006.2:p.Arg21Pro | missense |
| 11 | chr13:41240349 | T>C | *FOXO1* | 5,8/22,56 | 14.29 | NM_002015.3:c.1A>G | NP_002006.2:p.Met1? | start lost |
| 11 | chr16:11348989 | C>T | *SOCS1* | 45,45/202,271 | 15.99 | NM_003745.1:c.347G>A | NP_003736.1:p.Ser116Asn | missense |
| 11 | chr16:2983740 | C>G | *FLYWCH1* | 46,17/328,145 | 11.75 | NM_032296.2:c.1270C>G | NP_115672.2:p.Pro424Ala | missense |
| 11 | chr16:3788650 | T>A | *CREBBP* | 85,108/221,268 | 28.3 | NM_004380.2:c.4304A>T | NP_004371.2:p.Asp1435Val | missense |
| 11 | chr16:85954865 | G>T | *IRF8* | 46,57/290,294 | 14.99 | NM_002163.2:c.1258G>T | NP_002154.1:p.Glu420Ter | stop gain |
| 11 | chr17:40468887 | A>G | *STAT3* | 50,60/345,407 | 12.76 | NM_139276.2:c.2177T>C | NP_644805.1:p.Met726Thr | missense |
| 11 | chr22:23523770 | T>C | *BCR* | 45,41/336,392 | 10.57 | NM_004327.3:c.623T>C | NP_004318.3:p.Met208Thr | missense |
| 11 | chrX:66863138 | T>G | *AR* | 95,95/295,287 | 24.61 | NM_000044.3:c.1657T>G | NP_000035.2:p.Tyr553Asp | missense |
| 12 | chr12:57496661 | T>C | *STAT6* | 6,10/254, 274 | 2.94 | NM_003153.4:c.1256A>G | NP_003144.3:p.Asp419Gly | missense |
| 12 | chr13:41239875 | C>G | *FOXO1* | 2,9/54,141 | 5.34 | NM_002015.3:c.475G>C | NP_002006.2:p.Ala159Pro | missense |
| 12 | chr13:41240275 | C>T | *FOXO1* | 2,5/34,56 | 7.22 | NM_002015.3:c.75G>A | NP_002006.2:p.Trp25Ter | stop gain |
| 12 | chr16:11349033 | G>C | *SOCS1* | 8,9/86,145 | 6.85 | NM_003745.1:c.303C>G | NP_003736.1:p.Phe101Leu | missense |
| 12 | chr16:11349334 | A>G | *SOCS1* | 7,11/97,123 | 7.56 | NM_003745.1:c.2T>C | NP_003736.1:p.Met1? | start lost |
| 12 | chr16:3786695_3786697 | TCT>CC | *CREBBP* | 14,14/357,364 | 3.73 | NM_004380.2:c.4514_4516delinsGG | NP_004371.2:p.(Lys1505Argfs*45) | frameshift |
| 12 | chr16:3795276 | A>C | *CREBBP* | 14,8/260,149 | 5.1 | NM_004380.2:c.3914+2T>G | p.? | splice_donor_variant |
| 13 | chr1:2488146 | A>AC | *TNFRSF14* | 92,77/287,235 | 24.46 | NM_003820.2:c.48dup | NP_003811.2:p.Lys17GlnfsTer60 | frameshift |
| 13 | chr7:148508727 | T>G | *EZH2* | 80,40/388,199 | 16.97 | NM_004456.4:c.1937A>C | NP_004447.2:p.Tyr646Ser | missense |
| 13 | chr7:151945102 | G>A | *KMT2C* | 179,98/2644,2233 | 5.37 | NM_170606.2:c.2417C>T | NP_733751.2:p.Ser806Phe | missense |
| 13 | chr8:113697642 | AG>A | *CSMD3* | 91,35/412,148 | 18.37 | NM_198123.1:c.2474del | NP_937756.1:p.Thr825IlefsTer28 | frameshift |
| 13 | chr12:49423015 | C>A | *KMT2D* | 22,29/161,209 | 12.11 | NM_003482.3:c.14080G>T | NP_003473.3:p.Glu4694Ter | stop gain |
| 13 | chr12:49444932 | C>CG | *KMT2D* | 102,81/660,518 | 13.45 | NM_003482.3:c.2533dup | NP_003473.3:p.Arg845ProfsTer3 | frameshift |
| 13 | chr12:57498345 | C>T | *STAT6* | 19,31/101,115 | 18.8 | NM_003153.4:c.1114G>A | NP_003144.3:p.Glu372Lys | missense |
| 13 | chr15:45003780 | ACT>A | *B2M* | 37,33/194,206 | 14.89 | NM_004048.2:c.43_44del | NP_004039.1:p.Leu15PhefsTer41 | frameshift |
| 13 | chr16:3788617 | C>T | *CREBBP* | 130,161/241,236 | 37.89 | NM_004380.2:c.4337G>A | NP_004371.2:p.Arg1446His | missense |
| 13 | chrX:39913283 | T>C | *BCOR* | 44,94/254,376 | 17.97 | NM_017745.5:c.4730A>G | NP_060215.4:p.Glu1577Gly | missense |
| 14 | chr1:2491292 | C>T | *TNFRSF14* | 27,25/110,103 | 19.62 | NM_003820.2:c.335C>T | NP_003811.2:p.Ser112Phe | missense |
| 14 | chr1:27105692 | TTC>T | *ARID1A* | 23,31/178,188 | 12.86 | NM_006015.4:c.5305_5306del | NP_006006.3:p.Leu1769ArgfsTer3 | frameshift |
| 14 | chr2:128044582 | A>T | *ERCC3* | 25,45/160,204 | 16.13 | NM_001303416.1:c.847T>A | NP_001290345.1:p.Ser283Thr | missense |
| 14 | chr2:242076658 | C>A | *PASK* | 14,14/90,88 | 13.59 | NM_015148.3:c.898G>T | NP_055963.2:p.Val300Phe | missense |
| 14 | chr7:2976769 | C>T | *CARD11* | 41,36/165,142 | 20.05 | NM_032415.4:c.1243G>A | NP_115791.3:p.Asp415Asn | missense |
| 14 | chr7:2983864 | GCTC>G | *CARD11* | 27,25/180,131 | 14.33 | NM_032415.4:c.663_665del | NP_115791.3:p.Arg221del | inframe deletion |
| 14 | chr12:57496661 | T>C | *STAT6* | 48,43/225,210 | 17.3 | NM_003153.4:c.1256A>G | NP_003144.3:p.Asp419Gly | missense |
| 14 | chr16:3789597 | C>A | *CREBBP* | 53,45/137,103 | 28.99 | NM_004380.2:c.4262G>T | NP_004371.2:p.Cys1421Phe | missense |
| 15 | chr1:2489189 | G>GC | *TNFRSF14* | 79,82/175,208 | 29.6 | NM_003820.3:c.99dup | NP_003811.2:p.Cys34LeufsTer43 | frameshift |
| 15 | chr1:9777641 | A>AGCTCATCCAGGGCAGCAAAGT | *PIK3CD* | 12,4/159,137 | 5.13 | NM_005026.3:c.979_999dup | NP_005017.3:p.Leu327_Val333dup | inframe insertion |
| 15 | chr6:41903693 | TGTGACATCTGTAGGA>T | *CCND3* | 24,28/226,249 | 9.87 | NM_001136017.2:c.606_620del | NP_001129489.1:p.Pro203_Thr207del | inframe deletion |
| 15 | chr12:49420988 | A>AG | *KMT2D* | 60,69/293,328 | 17.2 | NM_004380.2:c.5039_5041del | NP_004371.2:p.Ser1680del | inframe deletion |
| 15 | chr12:57493839 | T>C | *STAT6* | 65,73/204,276 | 22.33 | NM_001429.3:c.2540dup | NP_001420.2:p.Ser848LysfsTer61 | frameshift |
| 15 | chr16:3781323 | AAGG>A | *CREBBP* | 107,121/158,137 | 43.59 | NM_005321.2:c.140C>G | NP_005312.1:p.Ala47Gly | missense |
| 15 | chr22:41545920 | T>TC | *EP300* | 72,84/329,401 | 17.61 | NM_170606.2:c.5053G>T | NP_733751.2:p.Ala1685Ser | missense |
| 16 | chr12:57498330 | C>T | *STAT6* | 16,22/50,67 | 24.52 | NM_003153.4:c.1129G>A | NP_003144.3:p.Glu377Lys | missense |
| 16 | chr13:41240288 | C>G | *FOXO1* | 16,26/37,52 | 32.06 | NM_002015.3:c.62G>C | NP_002006.2:p.Arg21Pro | missense |
| 16 | chr16:11349019 | C>T | *SOCS1* | 51,55/71,96 | 38.83 | NM_003745.1:c.317G>A | NP_003736.1:p.Ser106Asn | missense |
| 16 | chr16:3786123 | CATTGGGCCA>C | *CREBBP* | 15,11/61,55 | 18.31 | NM_004380.2:c.4633_4641del | NP_004371.2:p.Trp1545_Asn1547del | inframe deletion |
| 16 | chr16:85954886 | T>G | *IRF8* | 10,12/98,117 | 9.28 | NM_002163.2:c.1279T>G | NP_002154.1:p.Ter427GluextTer24 | stop lost |
| 16 | chrX:70613247 | C>T | *TAF1* | 27,27/75,78 | 26.09 | NM_004606.3:c.3208C>T | NP_004597.2:p.Arg1070Cys | missense |
| 16 | chr6:26156758 | C>G | *HIST1H1E* | 55,41/109,101 | 31.37 | NM_003482.3:c.14760dup | NP_003473.3:p.Ala4922GlyfsTer10 | frameshift |
