## Supplementary material for "*CREBBP* and *STAT6* co-mutation and 16p13 and 1p36 loss define the t(14;18)-negative diffuse variant of follicular lymphoma": Table 1

**Table 1. Patient demographic and pathologic characteristics.**

| **Case No.** | **1** | **2** | **3** | **4** | **5** | **6** | **7** | **8** | **9** | **10** | **11** | **12** | **13** | **14** | **15** | **16** | **Total** |
| --- | --- | --- | --- | --- | --- | --- | --- | --- | --- | --- | --- | --- | --- | --- | --- | --- | --- |
| **Age** | 45 | 57 | 64 | 38 | 51 | 44 | 46 | 38 | 59 | 48 | 56 | 66 | 66 | 67 | 50 | 56 | 53.2 (mean) |
| **Gender** | F | F | M | M | F | F | F | F | F | M | M | M | F | F | F | F | 5:11 (M:F) |
| **Morphology** | | | | | | | | | | | | | | | | |  |
| **Diffuse Growth (%)** | 100 | >75 | >75 | 100 | 100 | >75 | >75 | >75 | 75 | >75 | >75 | >75 | 100 | >75 | 100 | ~75 | 75-100% |
| **Micro-follicles** | - | + | + | - | - | + | + | - | + | - | + | - | - | + | - | - | 43.7% |
| **Sclerosis** | - | + | + | - | + | + | + | - | - | + | + | + | - | - | + | - | 56.2% |
| **Entrapped normal LN** | + | + | - | - | - | + | + | + | - | - | - | + | - | - | - | - | 37.5% |
| **Nuclear contour** | R | C | C | C | R | R&C | C | C | R | R | C | R&C | R | C | R | C | 5:4 (C:R) |
| **Immunophenotype** | | | | | | | | | | | | | | | | |  |
| **CD10** | + | + | + | + | + | + | +(P) | + | +(P)/W | + | + | + | + | + | + | + | 100.0% |
| **BCL6** | + | + | + | +(Fo) | NA | + | +(P) | +(Fo) | +(WP) | +(W) | + | + | + | + | + | NA | 100.0% |
| **BCL2** | + | +(V) | + | + | + | +(W)/E | E | E | +/- | + | + | + | + | + | - | + | 81.2% |
| **CD23** | +(P) | + | + | + | + | +(WP) | +(P) | +(Fo) | +/- | +(Fo) | + | + | + | +(W) | + | + | 100.0% |
| **FDC Network** | - | - | NA | - | - | - | Fo | Fo | - | - | Fo | - | - | Fo | - | Fo W | 31.2% |
| **Ki-67 Proliferation (%)** | <30 | 30-40 | 10 | <10 | NA | NA | 20 | NA | <10 | 10-20 | NA | 20-30 | NA | 10-20 | NA | 10-20 | 15-20% (mean) |

(Abbreviations. C: Cleaved; E: Equivocal; F: Female; Fo: Focal; FDC: Follicular Dendritic Cell; LN: Lymph Node; M: Male; NA: Not Available; P: Patchy; R: Round; V: Variable; W: Weak; _/_: When staining pattern of diffuse areas and micro-follicles differed, the “_/_” designation is used with the diffuse staining pattern on the left and the micro-follicle staining pattern on the right)
